# UMITIC: An unsupervised framework for the joint characterization of cellular phenotypes and spatial neighborhoods in multiplex and hyperplex immunofluorescence imaging data

**DOI:** 10.64898/2026.05.29.728633

**Authors:** María Sangüesa, Carlos E. De Andrea, Mikel Ariz

## Abstract

Multiplexed imaging technologies enable the simultaneous measurement of dozens of protein markers while preserving context, providing a view of tissue organization. However, extracting insights from these high-dimensional datasets—particularly in hyperplex settings (>20 markers)—remains a computational challenge, especially in the absence of annotated data. Here, we present UMITIC (Unsupervised Analysis of Multiplex Images via TIssue Characterization), a modular and unsupervised computational framework for the characterization of cell phenotypes and tissue neighborhoods from multiplex imaging data. UMITIC integrates three components: (i) CellCut, which combines nuclear and cytoplasmic predictions to improve cell delineation; (ii) CellMap, a contrastive learning approach that generates low-dimensional representations of image crops enriched with morphological features; and (iii) TissueNet, a graph neural network that models spatial cell–cell interactions to identify tissue neighborhoods. We evaluated UMITIC across four datasets to assess its robustness, scalability and biological relevance. In a 7-plex human tonsil dataset, UMITIC identified canonical immune cell populations and reconstructed well-established anatomical regions. In a 43-plex tonsil image, it preserved these structures while enabling a finer cell subtype stratification. In a 58-plex colorectal cancer cohort, UMITIC recovered previously reported immune composition differences and spatial organization variations between patient groups with different prognoses. Finally, on an expert-annotated mass cytometry imaging dataset of human lung tissue, UMITIC achieved higher agreement with reference tissue annotations than existing approaches, demonstrating improved lung microanatomy reconstruction. Together, these results show that UMITIC enables consistent and interpretable analyses of cellular phenotypes and tissue architectures across multiplex and hyperplex imaging datasets without manual annotations.

## Introduction

Advances in biomedical imaging and molecular profiling have enabled tissue analyses to be performed at unprecedented resolutions. The traditional single-cell profiling techniques, such as multiparameter flow cytometry or single-cell RNA sequencing, provide valuable molecular details but lack spatial context and morphological information. In contrast, high-dimensional technologies such as multiplexed imaging (MI) enable the simultaneous visualization of multiple biomarkers while preserving spatial information, thereby offering a more comprehensive view of tissue organization schemes [1], [2]. Recent technological advances have given rise to hyperplexed imaging approaches that are capable of simultaneously detecting hundreds of protein markers within a single tissue section, far exceeding the capacity of earlier multiplexed platforms. However, extracting biologically meaningful information from such complex data remains a major challenge in translational research [3], [4], [5].

This challenge is particularly evident in studies involving tumors, which are complex and heterogeneous cellular ecosystems wherein malignant, stromal and immune cells coexist and interact. Understanding the cellular compositions and spatial organization layouts of tumors is key to deciphering their behavior and progression [6]. Tissue architectures, combined with the expressions of predictive biomarkers, play a crucial role in tumor classification tasks and in guiding therapeutic decisions, especially in the context of immunotherapy [7]. Characterizing both cellular types and neighborhoods is essential for understanding tissue biology and identifying clinically relevant features, as spatial interactions capture the microenvironmental context beyond cell type identities and drive key immunological processes such as migration, antigen presentation, and cytotoxicity [8], [9].

Deep learning has emerged as a promising avenue for uncovering patterns in large-scale tissue imaging datasets [10]. However, most of the current approaches rely on supervised learning and require extensive manual annotations, which are time-consuming and difficult to scale across numerous biomarker panels and tissue types [11]. In the MI domain, the use of multiple biomarkers further increases the complexity of the annotation process [12], [13]. In this scenario, unsupervised methods offer compelling alternatives that enable the extraction of biologically meaningful insights without annotated data [14], [15]. Recently, deep learning frameworks have demonstrated success in spatial domain identification within the spatial omics field. For instance, recent methods such as stSCI [16], DisConST [17] and stMSA [18] have explored multi-task learning, distribution-aware contrastive learning and spatial multi-slice integration, respectively, to improve the identification and characterization of spatial domains. Despite these methodological advances in transcriptomics, jointly resolving single-cell morphological, spatial and phenotypic features in highly multiplexed protein imaging has not yet been fully addressed.

Recent MI analysis efforts have produced computational methods for studying the compositions and architectures of tissues. However, while methods such as UTAG [19], which focuses on the identification of cellular neighborhoods through graph-based local spatial interaction modeling, Naronet [20], a patch-based framework that learns clinically relevant tissue phenotypes, cellular neighborhoods and their interactions from highly multiplexed imaging data, and CytoCommunity [10], which identifies cellular neighborhoods from predefined cell phenotypes and their spatial distributions using graph neural networks, have advanced the MI analysis field, no such method simultaneously enables single-cell resolution phenotyping, spatial neighborhood identification, and direct interpretability, limiting the applicability of these techniques in scenarios with high-dimensional MI data [21].

To address these limitations, we introduce UMITIC (Unsupervised Analysis of Multiplex Images via TIssue Characterization), a fully unsupervised modular framework for conducting cellular-level tissue characterization in multiplex immunofluorescence imaging cases (Fig. 1). We first identify the nuclear and membrane boundaries of each cell using CellCut, enabling an accurate nucleus–cell pairing process and the reliable extraction of both morphological and marker localization features at a single-cell resolution. We then generate label-free cellular representations using CellMap, a contrastive self-supervised learning framework that encodes single-cell images into compact embeddings while reducing their dimensionality and mitigating technical variability, staining noise and intersample heterogeneity. The resulting embeddings are further enriched with morphological features and spatial coordinates. Cell-to-cell spatial interactions are modeled using TissueNet, a lightweight graph neural network that is developed in this work to learn context-aware cellular representations and capture biologically meaningful tissue neighborhoods without predefined spatial assumptions. This hierarchical design enables joint modeling of cellular phenotypes and spatial organization, leading to an interpretable tissue architecture reconstruction procedure at a single-cell resolution.

**Fig. 1.**
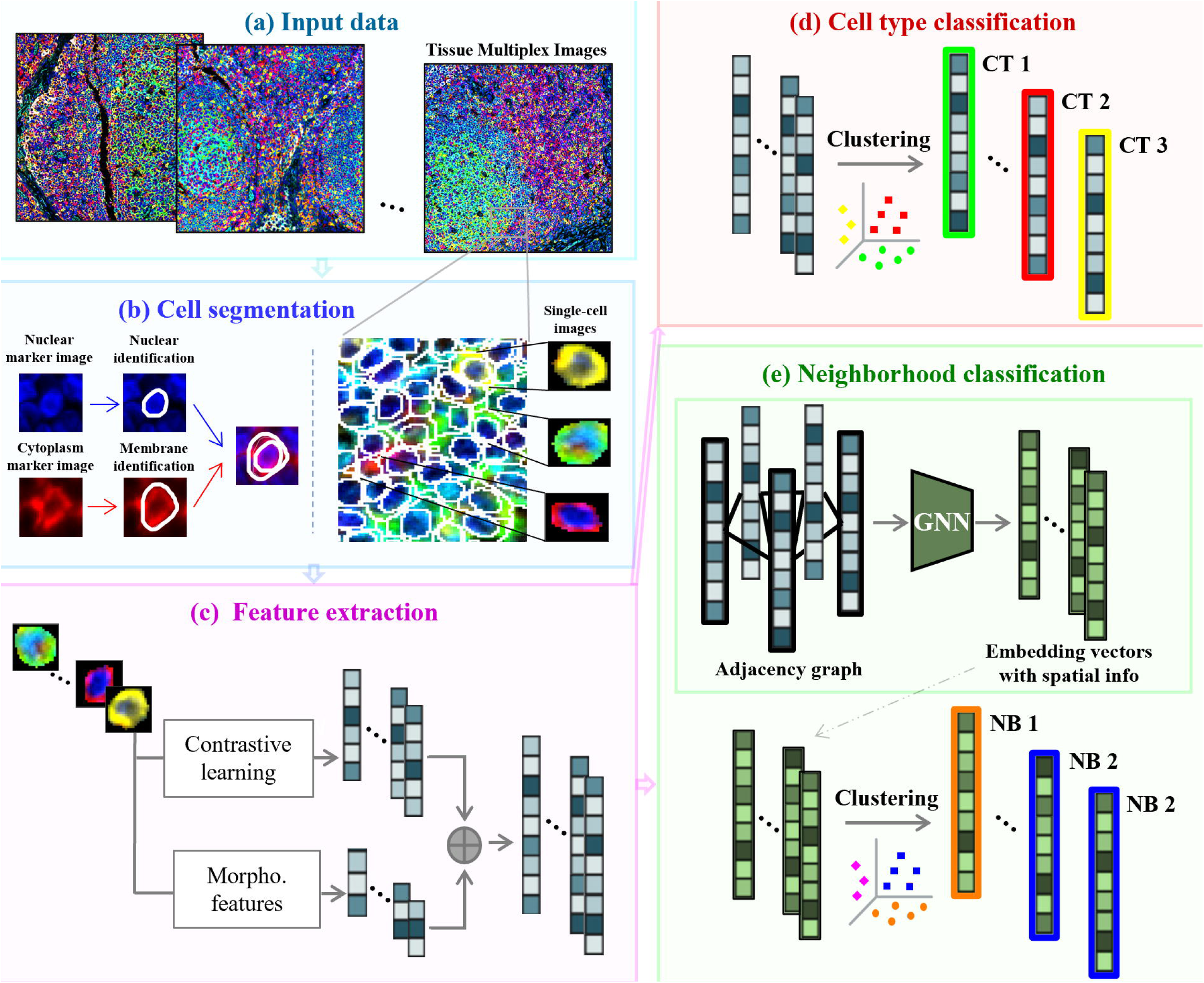
Overview of the proposed pipeline. **a.** Multiplex tissue images serve as inputs. **b**. A segmentation module generates nuclear and cytoplasmic masks, from which single-cell image crops are extracted. **c.** The single-cell images are embedded into low-dimensional vectors via a convolutional neural network trained with contrastive learning. Morphological features are added to enrich these embeddings. **d.** The cells are clustered on the basis of their vectors to identify cell types, assuming that similar embeddings correspond to the same type. **e.** An adjacency graph captures the spatial relations between neighboring cells. A graph neural network processes this graph to produce context-aware embeddings. The cells are then clustered (by considering both their own features and those of their adjacent cells) to identify spatial neighborhoods.

In addition to the conceptual novelty of a fully unsupervised and modular framework for conducting joint phenotypic and architectural tissue characterization, UMITIC introduces several methodological advances. Our main contributions include i) CellCut, a dual-segmentation strategy that takes advantage of two referenced state-of-the-art cell segmentation methods and postprocesses their segmentations to provide final, accurate, nucleus–cytoplasm paired single-cell delineation results; ii) CellMap, a contrastive self-supervised representation learning framework that is tailored to high-dimensional multiplex single-cell imaging scenarios, enabling robust and compact cellular embeddings that capture phenotypic and morphological information while mitigating technical variability and staining noise; iii) the use of the obtained accurate nucleus and cytoplasm segmentations to enrich the single-cell embeddings generated by CellMap with precise nuclear and cytoplasmic morphology descriptors, enhancing the ability to potentially discover functional or pathological disorders and providing more faithful representations of individual cells relative to those yielded by patch-based approaches; iv) TissueNet, a novel lightweight graph neural network that is designed to model spatial cell–cell interactions and infer context-aware tissue neighborhoods in a fully data-driven manner; v) a hierarchical and highly interpretable tissue architecture modeling strategy that jointly resolves cellular phenotypes and spatial neighborhoods at a single-cell resolution; and vi) an extensive experimental validation conducted across multiple independent multiplex and hyperplex imaging datasets, including healthy and pathological tissues, demonstrating that UMITIC consistently recovers biologically meaningful tissue organization schemes in different scenarios and outperforms state-of-the-art methods such as UTAG, CytoCommunity and NaroNet in both qualitative and quantitative analyses.

## Materials and Methods

Our methodology consists of four main steps. First (Fig. 1b), individual cells are segmented from tissue images, with nuclei and membranes explicitly identified. Second (Fig. 1c), each segmented cell is transformed into a low-dimensional feature vector using a contrastive learning-based approach, and this vector is enriched with cell morphology features. Third and fourth (Figs. 1d and e, respectively), each cell is associated with a phenotypic identity and assigned to a spatial neighborhood, respectively. Cell types are defined by marker expressions and morphologies, whereas neighborhoods are determined based on the spatial interactions between cells, incorporating both their phenotypic profiles and spatial contexts.

### Input Data

Three different datasets were used to assess the proposed approach: an in-house dataset consisting of 7-plex and 43-plex healthy tonsil tissue images, a public 28-plex Imaging Mass Cytometry (IMC) dataset including healthy lung tissue [22], and a 58-plex dataset containing colorectal cancer images [23].

The in-house dataset used to develop and evaluate our method consisted of multiplex (7 markers) and hyperplex (43 markers) immunofluorescence (IF) images of healthy human tonsil tissue. Tonsil tissue was used because of its well-characterized histology, facilitating the algorithm validation process [24], [25], [26]. Seven-plex images were generated using a multiplexed IF assay combining CD4, CD8, CD20, CD21, CD23, CK, and DAPI markers at the Department of Pathology at the Clínica Universidad de Navarra. The multiplexed IF assay development and validation procedures have been previously described by [27]. All tissues were used after receiving approval from the University of Navarra Human Research Committee (protocol number: 2022.109). The 7-plex dataset included six whole-slide images with sizes of 6000 × 6000 × 7 (pixels × pixels × markers). The 43-plex antibody panel and staining protocol were designed and validated using PhenoImager-Fusion 2.3.1 from Akoya Biosciences, a Quanterix Company. This panel comprised protein markers spanning numerous cell types, including epithelial cells, immune populations, structural features, and immune activation markers. Each image consisted of a single whole-slide human tonsil with dimensions of 9000 × 12000 × 43 (pixels × pixels × markers). The antibody panel contained the following proteins: DAPI-01, HLA-A, CD20, vimentin, CD31, SMA, PanCK, CD34, CD14, CD8, E-cad, CD45RO, CD44, GranB, iNOS, LAG3, HLA-E, CD19, Ki67, TIM3, Pax5, TIGIT, IFNG, VISTA, CD21, CD4, CD68, CD11c, CD45, CD3e, HLA-DR, CoIIV, FoxP3, CD163, IDO1, CD11b, CD57, PD1, PD-L1, CD107a, CD40, CD56 and CD278.

Unlike other imaging methods, such as hematoxylin and eosin (H&E) staining, whose algorithmic performance can be benchmarked using large public datasets [28], multiplex and hyperplex imaging techniques currently lack open, cell-level annotated datasets for conducting objective evaluations. To address this gap, we employed a publicly accessible, expert-annotated IMC dataset to objectively assess the ability of our pipeline to identify tissue architecture patterns [22]. This dataset comprised 26 highly multiplexed IMC images of healthy human lung tissue obtained from three donors, each encompassing 28 protein markers. Expert pulmonary pathologists manually annotated each image to delineate their organ-specific microanatomical domains, including airways, airway walls, submucosal glands, blood vessels, alveolar spaces, and cartilage.

Finally, to assess the generalizability of UMITIC to a clinically relevant, high-dimensional external dataset, we applied the full pipeline to a published 58-plex CODEX colorectal cancer cohort [23], comprising 140 tissue region images with sizes of 960 × 720 × 58 (pixels × pixels × markers) acquired from 35 patients. In this dataset, the patients were stratified into Crohn’s-like reaction (CLR) or diffuse inflammatory infiltration (DII) groups based on their spatial organization schemes and immune tumor microenvironment compositions. CLR tumors are characterized by the presence of organized tertiary lymphoid structures (TLSs) and are generally associated with favorable prognoses, whereas DII tumors exhibit more diffuse and less structured immune infiltration schemes and are linked to poorer clinical outcomes.

### CellCut: Cellular Segmentation (Fig. 1b)

We implemented CellCut, a hybrid cell segmentation model that produces accurate and consistent delineations of both nuclear and cytoplasmic regions. CellCut builds upon two complementary state-of-the-art deep learning components, StarDist [29] for nuclear detection purposes and DeepCell [30] for whole-cell contour estimation, but integrates them within a novel, unified segmentation workflow that is specifically designed to achieve enhanced nuclear–cytoplasmic pairing and boundary precision. For attaining improved accuracy, the CNN-based StarDist model was fine-tuned on a manually annotated in-house dataset consisting of 10,000 nuclei labeled from various multiplex images of different human tissue types. Nuclei whose detection probabilities were less than 60% were discarded. DeepCell was applied without fine-tuning because of the lack of an extensive annotated whole-cell dataset. A limited set of manually annotated whole-cell images was used to evaluate CellCut performance. The input images consisted of i) a nuclear signal for the nuclear model and ii) a cytoplasmic or membrane signal for the whole-cell model. In our case, DAPI served as the nuclear input, while the sum of cytoplasmic fluorescent markers, followed by anisotropic filtering, was used as the input for the whole-cell segmentation task, given the absence of a universal membrane marker.

To integrate both outputs for generating high-precision whole-cell masks, thereby addressing inconsistencies in which the nuclear and whole-cell segmentations did not identify a matching nucleus and cytoplasm for each cell, we included a postprocessing step in CellCut that ensured a one-to-one correspondence between each nucleus and a single associated cytoplasm. Conflict cases in which either one nucleus overlapped with multiple cytoplasms or one cytoplasm contained more than one nucleus were resolved using a watershed-based correction technique. Specifically, we constructed a binary mask containing only the conflicting nuclear and cytoplasmic regions. The watershed seeds were defined as the eroded nuclei involved in the conflict, while the grayscale input image for the algorithm was constructed by summing the intensities of all cytoplasmic markers and performing smoothing with anisotropic diffusion to enhance the continuity of the membrane signals. This approach allowed for the accurate redistribution of cytoplasmic boundaries around the nuclei, resulting in consistent and anatomically feasible whole-cell masks. Cells lacking either a nucleus or a cytoplasm—those identified by only one of the models—were excluded to retain only complete, analyzable cells. Cells without nuclei could correspond to red blood cells, which are typically removed prior to conducting image analyses because of their limited relevance in immune profiling cases [31], or to tangentially cut cells, which are excluded because the absence of the nucleus prevents the assessment of nuclear morphology and nuclear-localized protein expression. Fig. 2 shows examples where our approach corrected whole-cell segmentation errors made by DeepCell.

**Fig. 2.**
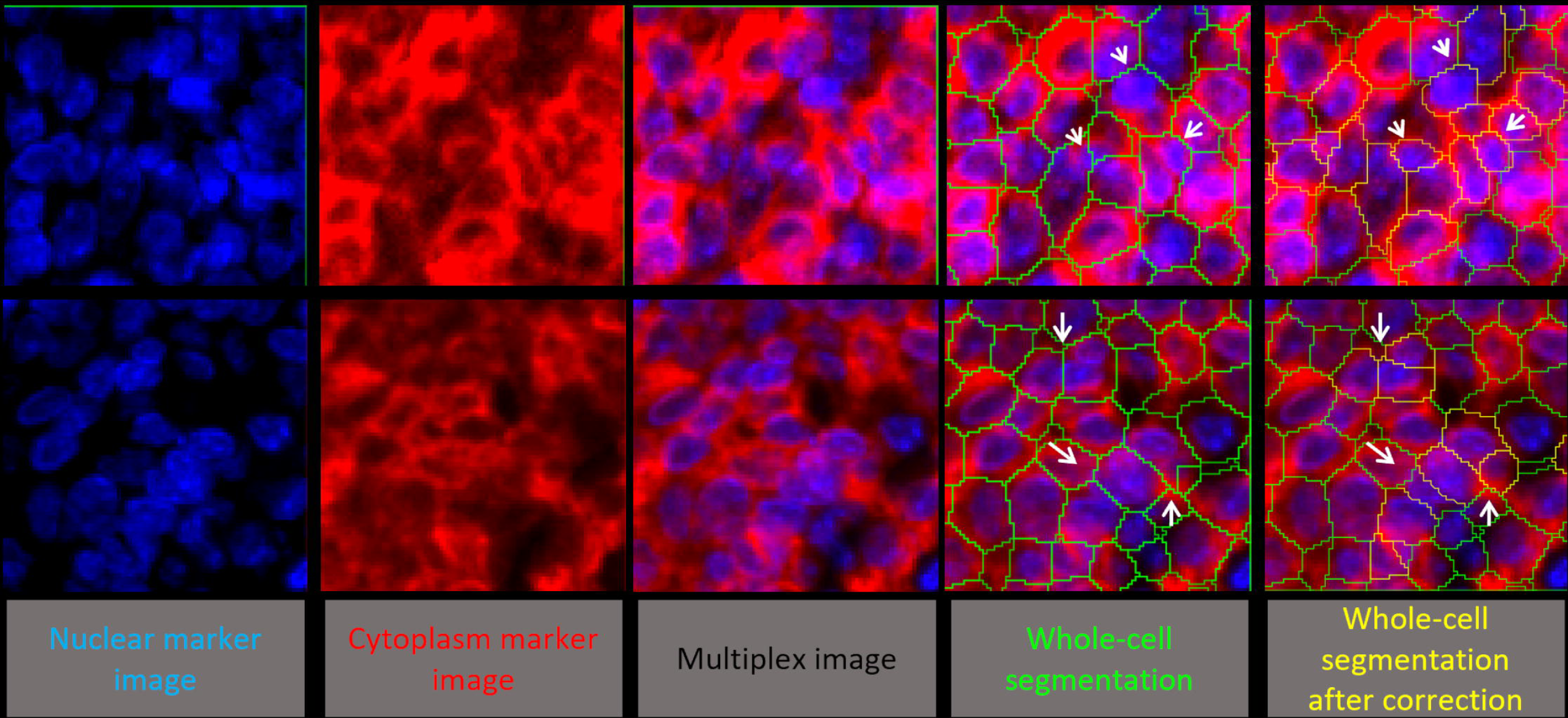
The corrected whole-cell segmentation process. Each row corresponds to a different sample. **a.** Nuclear marker image. **b.** Cytoplasmic marker image. **c.** Composite multiplex image combining nuclear and cytoplasmic channels. **d.** Segmentation result yielded by DeepCell, showing the initial cell boundaries. **e.** Whole-cell segmentation results obtained after performing correction with CellCut (yellow outlines). Arrows indicate segmentation errors: missing cytoplasm assignments for some nuclei, missed nuclei or a single nucleus split across multiple cytoplasmic regions. Our method corrects these errors, producing accurate one-to-one nucleus–cell assignments.

This dual-segmentation strategy enabled the accurate extraction of both nuclear and cytoplasmic regions. Each final whole-cell mask was used to extract a multichannel crop centered on each cell (Fig. 1b). These crops were then resized to a fixed size of, where was the predefined crop size and was the number of spectral channels of the input dataset. This resizing scheme ensured that the input requirements of the subsequent neural network were satisfied while preserving the essential cellular features by interpolating smaller cells and downsampling larger cells. Morphological cell features were calculated prior to this resizing operation.

### CellMap: Cell Feature Extraction (Fig. 1c)

To extract biologically meaningful representations from each multichannel single-cell image, we developed CellMap, a contrastive self-supervised learning approach that maps each cell to a low-dimensional embedding vector capturing spectral information (Fig. 3). Randomly sampled single-cell images (Fig. 3a) were augmented through random rotations and flips, channel-wise intensity variations, pixel dropout operations, affine transformations and Gaussian noise (Fig. 3b). Channel-wise intensity variations were limited to small multiplicative (0.9–1.1) and additive (±0.02) changes to mimic acquisition-and staining-related variability while preserving the relative marker-expression patterns that define cellular phenotypes. Both the original and augmented views were passed through a convolutional neural network based on a ResNet-50 backbone pretrained on ImageNet (Fig. 3c), which extracted high-level visual features. These features were then projected onto a 128-dimensional embedding space via a multilayer perceptron (MLP) (Fig. 3d). The models were trained using a contrastive loss function that combined the embeddings of positive pairs (original and augmented views of the same cell) while pushing the embeddings of negative pairs (different cells) apart. The loss for a positive pair was defined as follows:

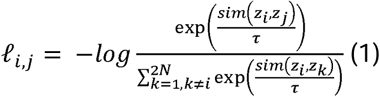

**Fig. 3.**
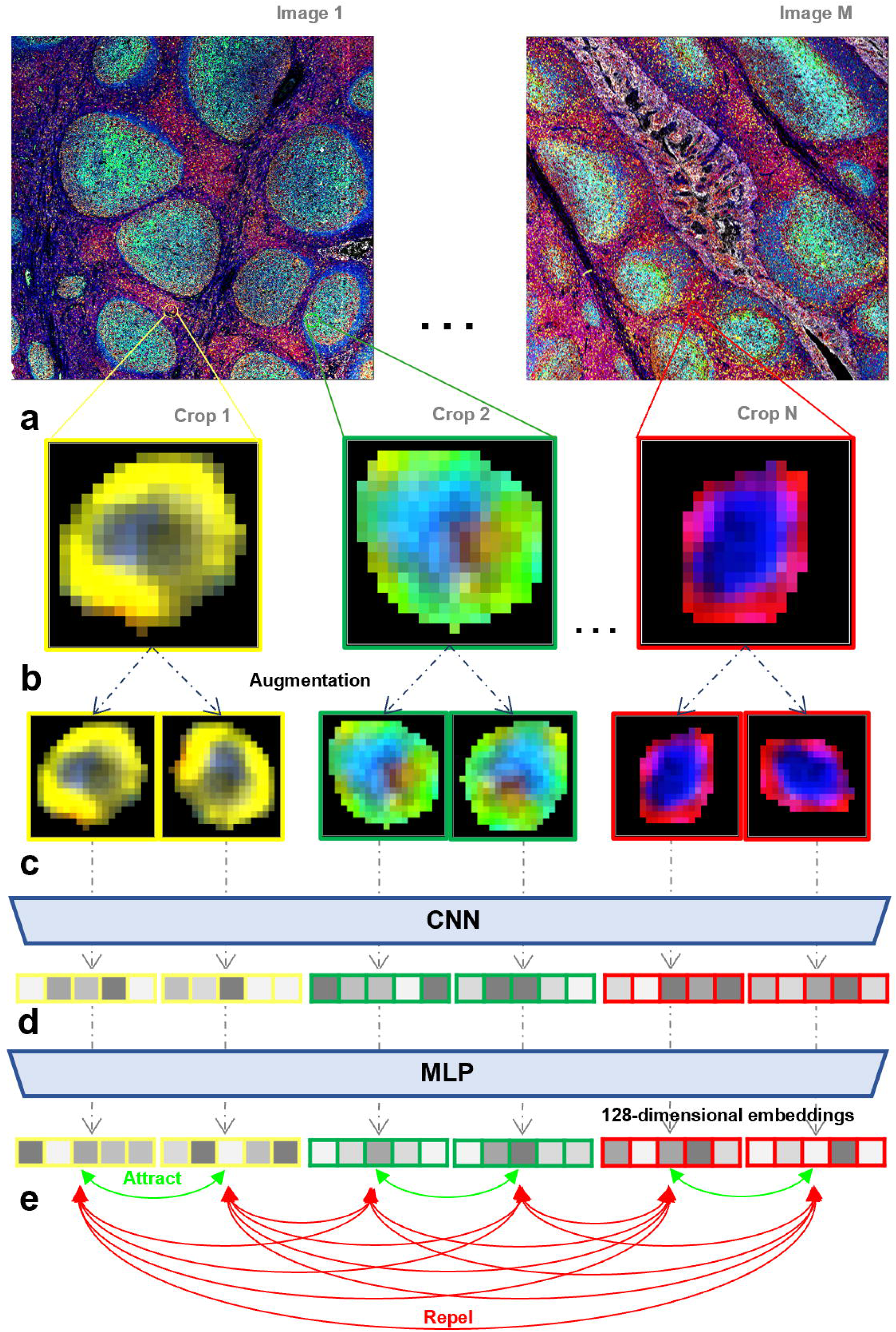
Iterative process of the automatic feature extraction module. Single-cell images **(a)** are input into the module, and augmented pairs are created for each cell **(b)** and passed into a backbone CNN **(c)** that learns high-level features, which are projected by an MLP **(d)** into a latent space where a contrastive loss pulls the representations of augmented views of the same cell together while pushing those of different cells apart **(e)**.

where τ is a temperature parameter (set to 0.07 in our case), *sim*(*u,v*) denotes the cosine similarity between vectors *u* and *v,z_i_*, and *z*_j_ correspond to the embeddings of a positive pair and *z*_k_ represents the embeddings of negative samples. Note that is the training batch size, resulting in a total of 2 views after applying the augmentation method. The training procedure iterated sampling, augmentation, feature extraction, projection, and parameter update steps via the loss function. This process enabled the learning of robust, manageable, biologically relevant representations of single-cell crops without manual annotations while reducing sensitivity to minor technical variability across acquisitions. One known challenge in MI is the variability and inter-and intrasubject differences that are introduced during the staining process. The above procedure is essential for reducing such noise, as the model learns to filter it out when patterns are found across different subjects.

To further enrich these embeddings, CellMap incorporated the morphological features extracted during the segmentation procedure (Fig. 1c), including i) cell areas in pixels; ii) eccentricity (0 = round, 1 = elongated); iii) cell diameters in pixels (estimated via principal component analysis as the greatest extents along the main axis); iv) nucleus-to-cell size ratios (0 = no nucleus, 1 = no cytoplasm); v) membrane shape complexity, computed as 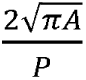 (*A* = area, *P*= perimeter), bounded in (0,1) with lower values meaning more irregular/elongated and higher values more regular/round membrane shapes; vi) cell polarity (0 = centered nucleus, 1 = nucleus touching membrane); and vii) nucleus areas in pixels. Accurately characterizing cellular morphologies is essential, as these features may be linked to underlying pathological processes [32], [33]. The resulting concatenated vector was z-score normalized.

### Cell Type Classification (Fig. 1d)

The embeddings resulting from CellMap captured both the morphological and marker expression features of each cell. We applied the Leiden clustering algorithm [34] to group cells into cellular categories in an unsupervised manner (Fig. 1d). The number of clusters was automatically determined on the basis of a user-defined resolution parameter, which controlled the coarseness of the clustering process and was the only required key input. This parameter can be adjusted depending on the biological question at hand, considering that higher resolutions will return more detailed subgroups, while lower values will produce coarser categories. Here, we empirically selected a resolution that reflected the well-characterized tonsil architecture to validate our method.

To reduce the imposed computational load, clustering was conducted in two stages. First, we randomly selected a subset of the cell embedding vectors and used them to define the clusters (i.e., the cell types) through the Leiden algorithm. Afterward, each remaining cell was assigned to the most frequently represented cell type among its nearest labeled cells in the embedding space. Note that the cell types were defined based on feature vectors derived from different images, ensuring consistent cell types between the subjects and enabling the implementation of comparative analyses.

### TissueNet: Spatial Context-Aware Cell Representation Learning via Graph Neural Networks (Fig. 1e)

To capture the spatial organization schemes of cells within tissues, we developed TissueNet, a self-supervised graph neural network (GNN) framework that learns context-aware cell representations to encode local tissue architectures. TissueNet identifies cellular neighborhoods—spatial tissue regions that are characterized by specific combinations of cell types. Unlike cell-centric approaches that rely solely on physical proximity or fixed cellular densities [35], [36], TissueNet dynamically identifies neighborhoods without requiring prior assumptions about their sizes or densities. The GNN operates over all cells within the tissue, integrating both cellular features and adjacency relationships to delineate meaningful neighborhoods, enabling a more flexible and generalizable analysis of tissue organization schemes.

To model the spatial relationships among cells, TissueNet builds an adjacency graph representing the tissue organization layout. This graph is defined by an adjacency matrix, where each cell is connected to its spatially closest neighboring cells if their center-to-center distance is at most twice the equivalent diameter target cell, which is estimated as the largest extent of the object along its principal axes using principal component analysis. The graph is denoted as *G=(V,E)*, where represents the vertices (cells) containing the embeddings of each image, and *E∈0,1M+M* represents the edges, which define the connectivity between vertices. Here, denotes the total number of cells contained in the tissue, which corresponds to the number of vertices in the graph. To reduce the memory requirements associated with storing full adjacency matrices, we represented the graph using a sparse adjacency matrix, storing only the nonzero edges in the edge list.

Our GNN leverages both cell embeddings and the associated spatial structure to update each cell representation in a more context-aware and discriminative manner by identifying the most relevant cell-to-cell interactions. Specifically, TissueNet implements a three-layer GraphSAGE architecture, in which each layer aggregates neighbor features by computing a mean over the sampled neighborhoods and concatenating the result with the representation of the target node, followed by a linear projection. The first layer projects the input embeddings into a 128-dimensional hidden space, the second layer applies a further nonlinear transformation in the same space, and the third layer produces the final 64-dimensional context-aware cell representations. ReLU activation functions and dropout are applied after the first two layers. We trained the model using a self-supervised contrastive learning paradigm. The training objective was to teach the model to distinguish between spatially adjacent node pairs (positives) and distant pairs (negatives). To achieve this, we implemented an InfoNCE loss function with an in-batch negative sampling strategy, as formulated in Equation 2. Within this scheme, for each connected pair of nodes (*i,j*), all the other nodes k served as negatives, without requiring explicit negative edge sampling. Specifically, a full cosine similarity matrix was computed over all node embeddings in each minibatch, and for each positive edge (i*,j*), the corresponding loss was computed as the negative log-softmax of the positive similarity against all other similarities in row.

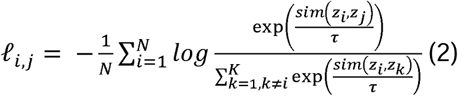

where z*_i_* is the embedding of a given cell, *z_j_* is the embedding of a positive sample (i.e., a spatially adjacent cell), _z*k*_ represents the embedding vector of a negative sample (i.e., spatially distant cells), *sim*(*u, v*) denotes the cosine similarity between vectors *u* and *v, τ* is a temperature parameter (set to 0.07 in our case) and K corresponds to the minibatch size minus one, since all nodes in the minibatch except the anchor served as negatives.

This training scheme compelled the network to generate embeddings where cells belonging to the same local microenvironment were mapped closer together in the feature space than cells derived from distant regions were, thereby effectively encoding tissue architectures. Finally, to delineate the cellular neighborhood, the resulting vectors were clustered using the Leiden algorithm to assign each cell to a distinct cellular neighborhood, following the same two-step clustering procedure described in the previous section.

## Results

To evaluate the performance and scalability of our proposed pipeline, we conducted four experiments with the datasets described in Section 2.1. First, we validated its viability using an in-house dataset consisting of multiplex images of six tonsil samples with a 7-channel biomarker panel. This setting enabled the simultaneous visualization of all channels, thereby facilitating a direct interpretation of the results. Second, we demonstrated the scalability of the method on a whole-slide hyperplex image of a tonsil sample containing 43 channels. While the complexity of the hyperplex image limited the ability to perform a direct visual analysis and posed computational and analytical challenges, we demonstrate that our method was capable of capturing consistent biological patterns that were identified in the 7-plex analysis. Afterward, we validated our pipeline on a 58-plex colorectal cancer database by identifying the findings described in the original publication [37]. Finally, we used the expert-annotated IMC dataset of human lung tissue to enable an objective comparative analysis with a state-of-the-art reference method in terms of unsupervised tissue architecture reconstruction [19].

### Tonsil 7-Plex Image Dataset

CellCut was applied using the DAPI channel for nuclear detection purposes, and the sum of the remaining channels was applied for cytoplasmic delineation. Each resulting 7-channel single-cell image crop with a size of 20 × 20 × 7 (pixels × pixels × markers) was encoded into a 128-dimensional embedding using the CellMap feature extraction module, achieving a top-1 contrastive accuracy of 97.86%, which was defined as the proportion of cells for which the augmented views of the same cells were ranked as the most similar embeddings. This metric reflects the quality of the representations learned during training; their biological relevance was validated through the downstream cell type identification results presented below.

Unsupervised clustering of these embeddings, which were previously enriched with morphological descriptors, enabled the identification of eight cell types (T1–T8; Fig. 4a) using a Leiden resolution parameter of 0.3. To incorporate spatial context, we applied our TissueNet approach, producing 64-dimensional embedding vectors, and performed clustering to find five cellular neighborhoods (NB1–NB5; Fig. 4b) with a Leiden resolution parameter of 0.5. Both heatmaps were column-normalized, and clustering was performed across the entire dataset.

**Fig. 4.**
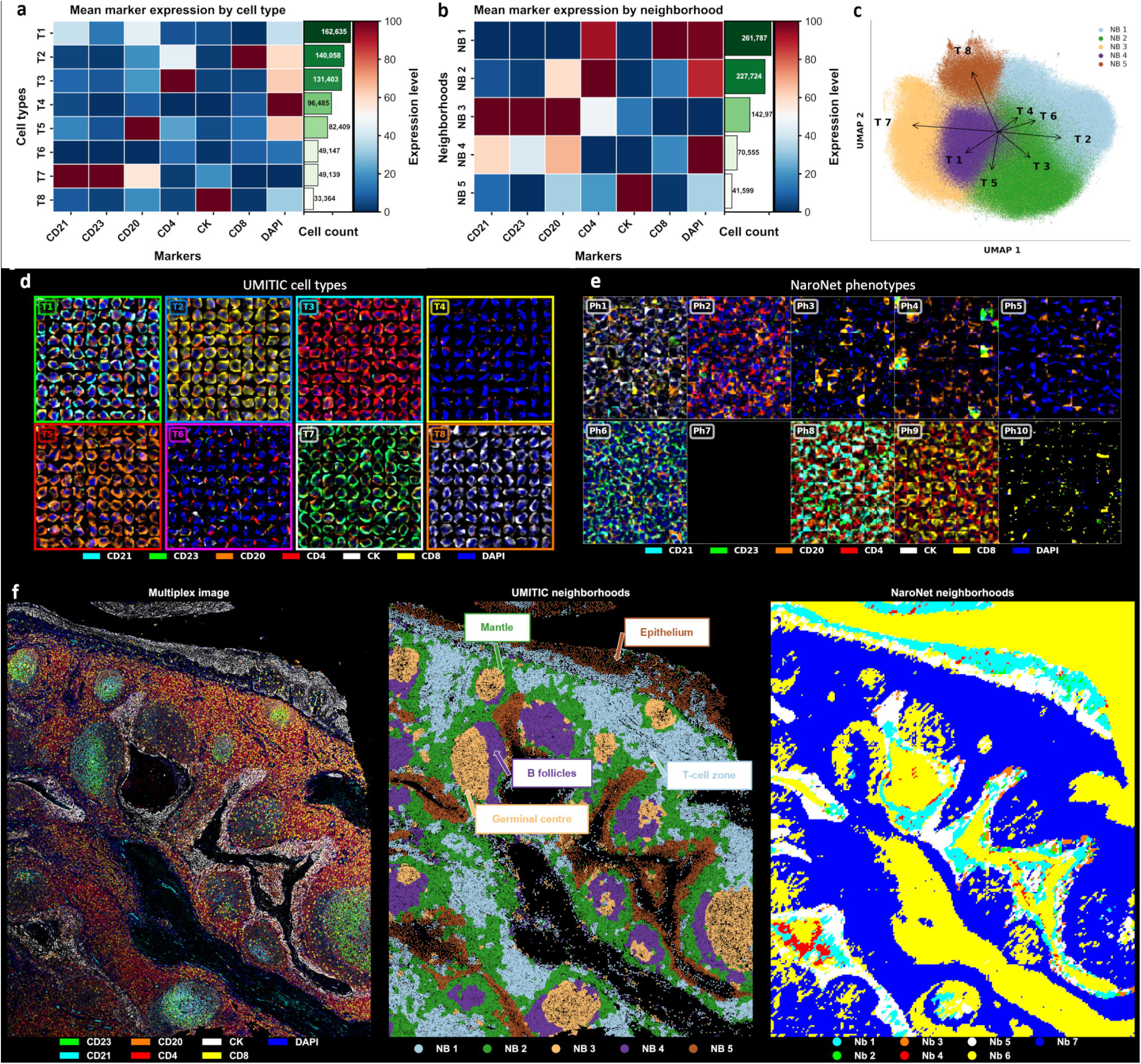
Results derived from six human tonsil samples analyzed with a 7-plex multiplex panel. **a**. Heatmap of the average marker expression per identified cell type (T1–T8), normalized by marker. The cell counts are shown in green. **b**. Heatmap of the average marker expression per neighborhood (NB1–NB5), normalized by marker. The cell counts are shown in green. **c.** UMAP of the cell embeddings generated by TissueNet. Each cell is colored according to its assigned neighborhood. The top four loadings of components 1 and 2 per cell type are shown as vectors. **d.** Mosaics of the UMITIC single-cell examples per identified cell type (T1– T8), showing consistent marker expression profiles. **e.** Mosaics of the NaroNet patch examples per identified cell type (Ph1–Ph10), showing consistent marker expression profiles. **f.** Comparison among the tissue reconstruction results (by neighborhood) obtained across the six tonsil samples at the baseline. Left: raw multiplex image; center: UMITIC reconstruction result; right: NaroNet reconstruction results. Known tonsillar regions—B follicles, germinal centers, mantle zones, T-cell zones, and epithelium—are annotated to associate them with the neighborhoods identified by UMITIC.

The CellMap module effectively encoded both marker expressions and morphological patterns, enabling the accurate and consistent identification of well-characterized cell types. The key cell populations, including CD20+CD21+ B cells (T1), CD8+ T cells (T2), CD4+ T cells (T3), negative cells (T4, T6), CD20+ B cells (T5), mature B cells characterized by CD20+CD21+CD23+ expressions (T7) and CK+ epithelial cells (T8), were reliably identified across the different samples. This alignment between the predicted cell types and the marker expressions is further supported by the heatmap shown in Fig. 4a. The consistency of the marker expression profiles at the single-cell level is highlighted in Fig. 4d via the display of the representative cells for each predicted cell type across the entire dataset. These examples demonstrate the ability of our approach to cluster phenotypically and morphologically similar cells regardless of their sample origins, reinforcing the robustness of the learned representations. Notably, the T4 and T6 cells share similar marker expression profiles (limited to DAPI staining – negative cells –) but differ in their morphologies: T4 cells are more elongated, with a mean eccentricity of 0.79 (compared to 0.51 in T6 cells). This illustrates the sensitivity of our pipeline to subtle morphological differences. Supplementary Figure 1a shows two different multiplex input images (top) along with their reconstructed versions based on the predicted cell type assignments (bottom).

The identified spatial neighborhoods corresponded closely to known tonsillar structures: T-cell zones (NB1), mantle zones (NB2), germinal centers (NB3), B follicles (NB4) and the epithelium (NB5) (Fig. 4f). The consistency of these neighborhoods across different samples is demonstrated in Supplementary Figure 1b, where similar spatially organized tonsillar regions were consistently recovered across independent tissue samples despite the presence of intersubject variability. The cell-type composition of each neighborhood (Supplementary Figure 1c) confirmed the meaningful assignations. T-cell zones (NB1) are composed mainly of CD8+ T cells (T2), CD4+ T cells (T3), and some negative cells (T4). Mantle zones (NB2) are composed primarily of CD4+ T cells (T3), CD20+CD21+ B cells (T1) and CD20+ B cells (T5). Mature B cells (T7) mainly appear at germinal centers (NB3). B follicles (NB4) are composed mainly of CD20+CD21+ B cells (T1). Finally, the epithelium (NB5) is composed primarily of CK+ epithelial cells (T8). These findings are perfectly consistent with the expected tonsil histology [25].

We projected the GNN-extracted cell embeddings into a two-dimensional space using UMAP [38], coloring each point by its neighborhood assignment to visualize the spatial clustering results (Fig. 4c). Cell type centroids were computed, and directional vectors from the global center of mass were added to emphasize which cell types (T1–T8) were most influential in each spatial cluster (NB1–NB5). For instance, NB3 (germinal centers) was positioned in the direction of the vectors corresponding to T1 (CD20+CD21+ B cells) and T7 (CD20+CD21+CD23+ B cells), indicating a close association between this neighborhood and B-cell populations; this is consistent with the known enrichment of mature B cells within germinal centers. In contrast, the direction corresponding to NB5 (epithelium) aligned exclusively with T8 (CK+ epithelial cells), highlighting the clear separation and distinct cellular composition of epithelial tissue.

To obtain a baseline comparison, we trained NaroNet in its unsupervised configuration on our 7-plex tonsil dataset. NaroNet [20] is a state-of-the-art weakly supervised deep learning framework that was originally developed for completing patient-level classification tasks by integrating multiscale tissue information (phenotypes, neighborhoods, and areas) from multiplex imaging data. It can also operate in a fully unsupervised mode, enabling exploratory tissue analyses without prior annotations. This comparison enabled a direct comparison between the patch-based paradigm (NaroNet) and our cell-based approach (UMITIC). In this setup, NaroNet inferred tissue phenotypes and neighborhoods from the input image tiles, with the patch size adjusted to approximate the mean cell area (20 pixels). The numbers of phenotypes and neighborhoods were set to 10 and 7, respectively, corresponding to the empirically obtained optimal configuration, since NaroNet requires these parameters to be manually set.

NaroNet identified ten phenotypes (Ph1–Ph10) that were conceptually analogous to the cell types defined in UMITIC (Supplementary Fig. 2a). However, its phenotype heatmap revealed mixed marker profiles, with biologically implausible combinations such as simultaneous CD8 and CD4 expressions (Ph9), originating from adjacent cells within the same patch. This occurred because each patch did not accurately represent a single cell. The representative example patches of the predicted phenotypes (Fig. 4e) exhibited consistent marker patterns; however, their interpretability remained limited, and the resulting phenotypes did not reflect the well-established cellular composition of tonsillar tissue.

Similarly, NaroNet identified seven neighborhoods (Nb1–Nb7) (Supplementary Fig. 2b). The baseline comparison in Fig. 4f benchmarks UMITIC against NaroNet in terms of tissue architecture reconstruction. This comparative analysis highlights the ability of UMITIC to resolve the tonsillar microanatomy, overcoming the limitations of NaroNet. While background and epithelial regions could be associated with neighborhoods Nb1 and Nb5 and Nb6, respectively, NaroNet failed to correctly distinguish between the T-and B-cell zones, with Nb7 displaying extensive overlap between these regions.

### Tonsil 43-Plex Image Dataset

To perform the CellCut step, the DAPI channel was used for nuclear identification purposes, and the sum of the CD20, PanCK and CD3e channels was used for cytoplasmic delineation. Each resulting 43-channel single-cell image crop consisting of 20 x 20 pixels was encoded into a 128-dimensional embedding, with 97.21% top1-contrastive accuracy reached using the CellMap module. These embeddings were further processed by our GNN to incorporate spatial context, producing 64-dimensional context-aware representations.

One key advantage of using a larger marker panel is the ability to perform finer analyses of cell types, with the potential to identify distinct cellular subtypes. To exploit this added granularity, which adds more detailed cell type assignments, we increased the resolution parameter for cell-type clustering to 2.5, which led to the identification of 56 distinct cell types (T1–T56) (Fig. 5a). At this resolution, all cellular families observed in the 7-plex experiment were preserved, while additional markers enabled further stratification into functional subpopulations. For instance, the T3 cells identified in the 7-plex panel, which corresponded to CD4⁺ T cells, were subdivided into multiple groups. These subgroups could be linked to known phenotypes, such as T5, which represents CD4⁺PD-1⁺ cells, and T17, which corresponds to CD4⁺FoxP3⁺ cells. High clustering resolutions are particularly useful for distinguishing cell types on the basis of variations in marker expression levels, especially in cases where a binary classification method with positive or negative marker expressions is insufficient. Functional markers such as PD1, GRANB or PD-L1 can be absent or expressed at low or high levels within a cell, and these expression levels often correlate with distinct functional states. For example, in our dataset, the T5 CD4⁺ T-cell subtype exhibited high PD-1 expression, whereas the T20 CD4⁺ T cells presented lower PD-1 levels, suggesting that T5 cells may be in a more activated or functionally distinct state than T20 cells are.

**Fig. 5.**
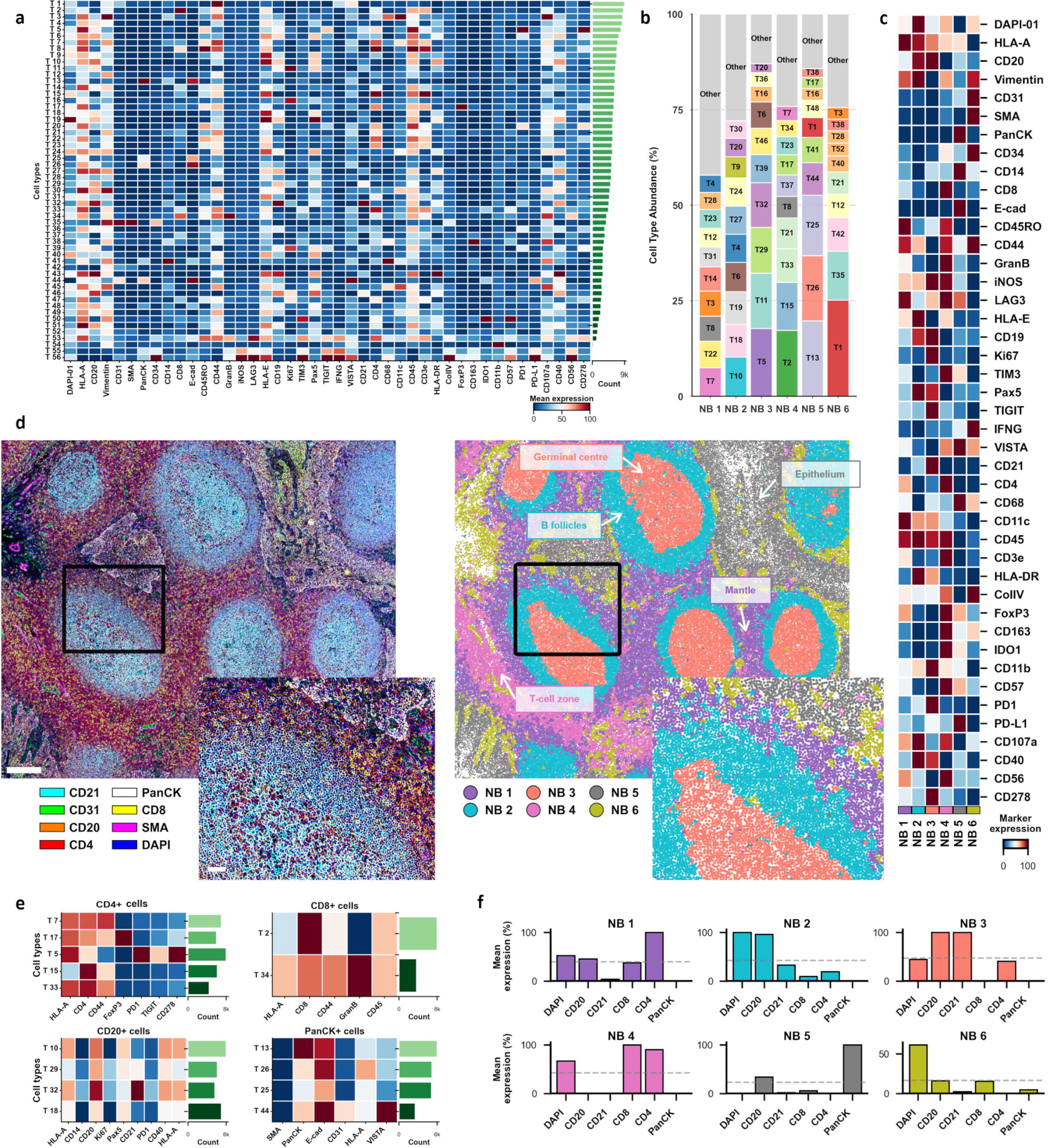
Results derived from the 43-plex tonsil image. **a.** Heatmap of the average marker expression per cell type (T1–T56; column-normalized). The cell counts are shown in green. **b.** Cell type abundances across different neighborhoods (NB1–NB6); only the 10 most abundant types per NB are shown for simplicity. **c.** Heatmap of the average marker expression per neighborhood (NB1–NB6; row-normalized). **d.** Tissue reconstruction outcome showing the original hyperplex image (left) and the neighborhood-based cell classification result provided by UMITIC (right), which is associated with well-known tonsillar regions. **e.** Marker expression heatmaps produced for representative subsets of the cell types, showing how the broader populations identified in the 7-plex experiment (CD4+, CD8+, CD20+, and PanCK+) are now subdivided into more refined phenotypes. **f.** Mean marker expression (extracted from Panel C) obtained for the subset of markers used in the 7-plex experiment, demonstrating that the same anatomical and cellular structures are recapitulated in the 43-plex data.

When the resolution was greater than 2.5, the differences between the additional clusters and those identified at 2.5 were minimal, mostly reflecting minor marker intensity variations that may lack biological relevance. In contrast, the employment of a lower resolution of 0.3, such as that used in the 7-plex experiment, merged several phenotypically distinct populations, identifying only 20 cell types and failing to capture functionally meaningful differences. Given that our current panel included 43 markers, a more detailed tissue composition characterization was needed. Importantly, the resolution parameter is user-configurable, enabling its adaptation to the desired granularity for specific research or clinical questions.

To facilitate the interpretation of the heatmap shown in Fig. 5a, we grouped the cell types by their key marker expressions (Fig. 5e). The main cellular families found in the 7-plex data, i.e., CD4-, CD8-, CD20-, and CK-expressing cells, were resolved into more specific cell subtypes, reflecting expression intensity differences caused by the higher resolution parameter and the contribution of the added functional markers, which provided information on the states and activities of the cells. These results illustrate the ability of the proposed method to achieve higher phenotypic resolutions as the marker dimensionality increases, providing deeper tissue composition insights.

In contrast, for the spatial neighborhood analysis, our goal was to assess whether the same tissue-level structures identified in the 7-plex data could also be recovered in this more complex setting. As such, we maintained the same resolution parameter (0.5) that was used previously since neighborhoods capture higher-order organization schemes rather than fine-grained cellular type diversity. With this setting, we identified six neighborhoods (NB1–NB6) across the entire tissue (Figs. 5c and 5d).

We then analyzed the composition of each neighborhood in terms of its cell type abundance, which revealed the ten most-frequent cell types per neighborhood (Fig. 5b). Utilizing these data, we associated the identified neighborhoods (Fig. 5c) with known tonsillar tissue compartments. To conduct a direct comparison with the 7-plex results, we focused on the markers that were shared by both panels (DAPI, CD20, CD21, CD4, CD8, and PanCK) to visualize the marker distributions across the identified neighborhoods (Fig. 5f). Note that the marker intensities were normalized for each marker to facilitate meaningful cross-neighborhood comparisons.

NB1 (mantle zones) was coexpressed with CD4, CD8, CD20, and CD21, which was consistent with the findings obtained in mixed T-and B-cell regions. NB2 (B follicles) and NB3 (germinal centers) were dominated by B-cell markers, with NB3 exhibiting higher CD21 expression levels than NB2 did, indicating cell maturation within the germinal centers. NB4 (T-cell zones) displayed high expression levels for both CD4 and CD8 markers. NB5 (epithelium) was enriched in PanCK-expressing epithelial cells. NB6 lacked significant expressions of the shared markers, suggesting that it corresponds to a tissue region defined by features that are unique to the 43-marker panel.

In summary, the 43-plex dataset demonstrates the scalability of our pipeline, enabling more detailed characterizations of cell phenotypes through a user-adjustable resolution while maintaining strong consistency in the detected tissue-level structures through stable neighborhood analyses.

### External CODEX 58-plex colorectal cancer cohort [23]

To validate the biological validity and robustness of UMITIC, we performed an external validation using the CODEX 58-plex colorectal cancer cohort that was previously analyzed in [37], which we used as a ground-truth reference.

During cell segmentation (CellCut step), the DRAQ5 channel was used for nuclear identification purposes, and the sum of the CD20, cytokeratin and CD3 channels was used for cytoplasmic delineation. Each resulting 58-channel single-cell image crop with 20 x 20 pixels was encoded into a 128-dimensional embedding, reaching 98.31% top1-contrastive accuracy after 100 epochs using the CellMap module. These embeddings were further processed by our GNN to incorporate spatial context, producing 64-dimensional context-aware representations.

Performing unsupervised clustering on the CellMap-extracted embeddings revealed 53 cell types using the 2.4 Leiden resolution, which was selected to match the reported level of cellular granularity, where the cell populations were originally identified through X-shift clustering [39] followed by manually guided merging. Biological identities were assigned to the resulting unsupervised clusters on the basis of their normalized marker expression profiles, following the interpretation strategy described in the original study. Supplementary Fig. 3 shows the marker expression heatmaps that were used to associate each cluster with its corresponding populations. Most of the major immune populations identified by UMITIC were consistent with those reported in the original study (Fig. 6a). Specifically, B cells, CD68+ macrophages, CD11b+CD68+ macrophages, and regulatory T cells were significantly more abundant in DII patients (p < 0.05–0.001, Mann–Whitney U test), whereas the numbers of NK cells, CD8+ T cells and other populations did not significantly differ between the groups, which was in agreement with the reported results.

**Fig. 6.**
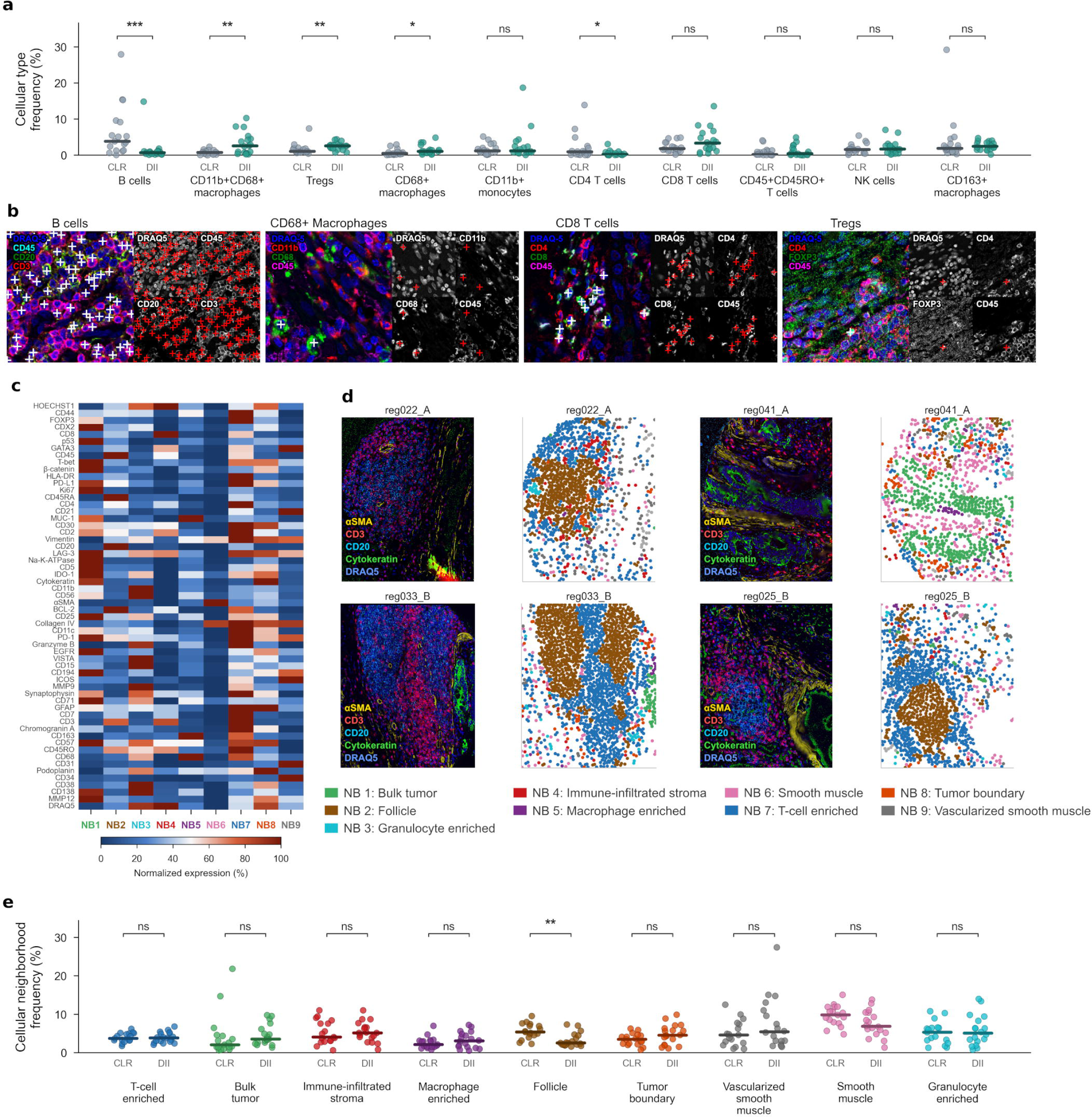
Validation of the cell type and neighborhood identification results obtained for the external 58-plex colorectal cancer dataset. **a.** Frequencies of immune cell populations across different patient subtypes. The significant differences among the abundances of B cells, macrophage subsets and regulatory T cells of CLR and DII patients is consistent with previously reported findings (* p < 0.1, ** p < 0.01, *** p < 0.001; Mann Whitney U test). **b.** Representative multichannel fluorescence images illustrating four immune cell phenotypes. For each phenotype, the composite overlay (left) and individual marker channels (right) are shown. The crosses indicate the centroids of the cells that are assigned to the corresponding phenotype described in the title. **c.** Heatmap showing the mean normalized expressions of 58 protein markers across the nine cellular neighborhoods. **d.** Cellular neighborhood reconstruction outcomes. For each region, its multichannel fluorescence image (left) and the corresponding neighborhood map (right) are shown, with each dot representing a single cell colored according to its assigned neighborhood. **e.** Frequency of each cellular neighborhood across the CLR and DII patient subtypes. Only the number of follicle neighborhoods (TLS) significantly differs between the patient groups (** p < 0.01, Mann Whitney U test), which is in agreement with previously published results.

The cell phenotype assignment results were further validated by marking cell centroids for the representative cells of different types and inspecting multichannel overlays and individual marker channels (Fig. 6b). For example, in a tissue region annotated for CD68⁺CD11b⁻ macrophages, the method correctly identified CD68-expressing cells lacking CD11b expressions. Similarly, B-cell populations were accurately identified on the basis of CD20 expressions, while CD3-expressing T lymphocytes were not assigned to these clusters, which was consistent with the expected marker expression profile and confirmed the agreement between the marker expressions and the assigned cell identities.

A neighborhood analysis was performed using a Leiden resolution of 0.7 to recover thecellular neighborhoods reported in the original study [37]. In the original work, these neighborhoods were identified by analyzing the abundance of the cell types contained within sliding tissue windows. Neighborhood identities were defined on the basis of their marker expression profiles and spatial localization statuses. This resulted in ten distinct neighborhoods, one of which was excluded from further analysis because of the presence of imaging artifacts, yielding nine biologically interpretable neighborhoods (Figs. 6c–d). Consistent with the original study, this artifact-associated neighborhood was excluded from the downstream biological analyses, further highlighting the ability of UMITIC to automatically separate nonbiological patterns from meaningful tissue organization schemes. The spatial tissue region reconstruction results demonstrated coherent and interpretable neighborhood maps at a single-cell resolution (Fig. 6d). For example, T-cell-enriched neighborhoods were observed surrounding follicular regions, reproducing the spatial organization of tertiary lymphoid structures described in the original study. Similarly, follicle-associated neighborhoods formed compact and well-delimited regions, which was consistent with the expected compartmented structures.

A comparative analysis of the neighborhood frequencies among different patient groups revealed significant follicular neighborhood enrichment, i.e., TLSs, in CLR patients relative to DII patients (p < 0.01, Mann Whitney U test), whereas no other neighborhoods showed significant differences (Fig. 6e). These findings are consistent with the main findings reported in the original paper: CLR tumors are characterized by the presence of organized TLSs, whereas these structures are absent in DII tumors.

Minor differences were nevertheless observed for some neighborhoods, such as smooth muscle and bulk tumor regions, whose relative abundances showed small shifts compared with those in the original study, despite remaining nonsignificant between the groups. These discrepancies may have stemmed from the different neighborhood definition strategies, as the original work relied on window-based spatial aggregation, whereas UMITIC assigns neighborhood identities at a single-cell resolution, enabling finer characterizations of local tissue organization structures.

### Annotated IMC dataset [22]

We benchmarked UMITIC against UTAG [19] and CytoCommunity [10], state-of-the-art reference methods for the unsupervised discovery of tissue architectures, using a publicly available, expert-annotated IMC dataset of lung samples [22]. Since the dataset provides only spatial neighborhood labels and UTAG and CytoCommunity cannot classify individual cell types, we specifically compared the spatial neighborhoods inferred by UMITIC (TissueNet module),UTAG and CytoCommunity against the available expert annotations (Fig. 7).

**Fig. 7.**
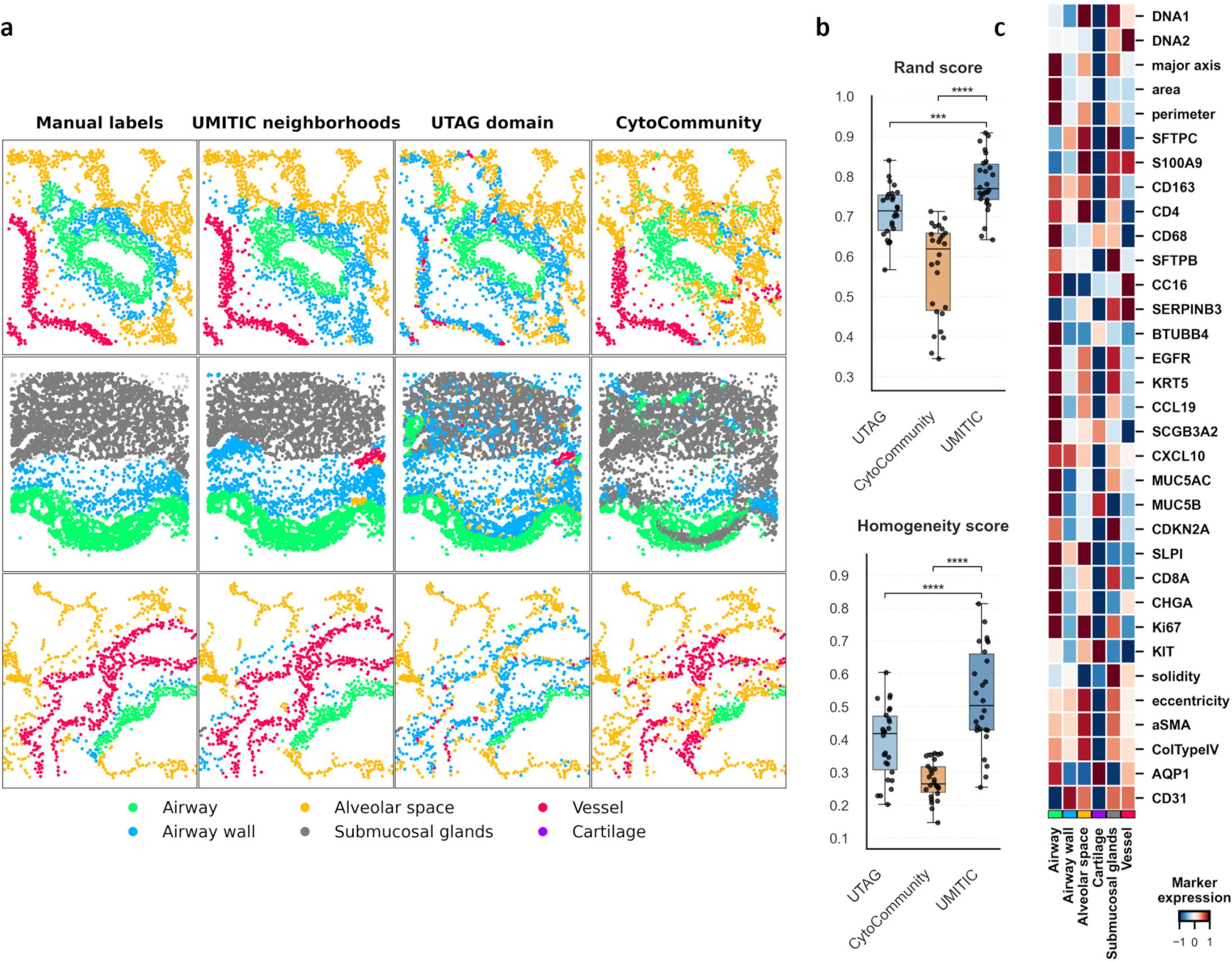
Comparative performance analysis between our GNN-based neighborhood inference process (the TissueNet module in UMITIC) and those of the state-of-the-art UTAG and CytoCommunity methods in terms of reconstructing the lung microanatomy. **a.** Representative IMC lung images annotated manually by pathologists (left) and processed by UMITIC (middle) and the state-of-the-art methods (right). **b.** Quantitative evaluation of the neighborhood inferences yielded by UMITIC, UTAG and CytoCommunity against expert annotations. Statistical significance was assessed by paired Wilcoxon signed-rank tests comparing, with p-values adjusted for multiple comparisons using the Benjamini–Hochberg method (***p < 0.001, ****p < 0.0001). **c.** Heatmap of the marker expression profiles produced across the tissue domains annotated in the dataset and retrieved by UMITIC in an unsupervised manner.

For this purpose, TissueNet was provided the single-cell spatial coordinates and marker intensity profiles that were available in the dataset. The learned spatial-aware embeddings were then clustered using the Leiden algorithm (with a resolution of 0.3) and produced distinct cellular neighborhoods (Fig. 7a), which were subsequently annotated on the basis of their marker expression profiles (Fig. 7c).

The neighborhood inference performance of the tested methods was quantified using the Rand index, which measures the degree of label concordance between the obtained neighborhoods and the expert annotations, and the homogeneity score, which evaluates how uniquely the predicted clusters correspond to expert annotations. Both are reference metrics for measuring accuracy in tissue domain characterization tasks and were reported in the original UTAG paper.. For benchmarking, UTAG was evaluated using the configuration reported in its original study for this dataset, with the same Leiden resolution used in UMITIC (0.3). CytoCommunity was run with 12 TCNs, which yielded the highest scores, with all remaining parameters set to their default values. UMITIC achieved higher homogeneity and Rand index scores than both methods (0.526 ± 0.153 and 0.782 ± 0.072, respectively), compared with 0.391 ± 0.110 and 0.710 ± 0.063 for UTAG and 0.259 ± 0.085 and 0.582 ± 0.104 for CytoCommunity (p < 0.001 for all comparisons). Statistical significance was assessed using paired Wilcoxon signed-rank tests.(Fig. 7b). A qualitative evaluation of the IMC samples shown in Fig. 7a further supports this statement. These findings underscore the strength of our GNN-based approach in terms of capturing biologically meaningful spatial domains within complex tissues.

### In-depth analysis

#### Comparison of CNN-based and marker-intensity representations

To assess the contribution of the learned CellMap representation beyond marker intensity alone, we compared our CNN-based embeddings with a baseline representation consisting of per-cell marker intensity vectors. In the 7-plex tonsil dataset, clustering based solely on marker intensities identified cell phenotypes with heterogeneous representation across samples, with some phenotypes being detected predominantly in individual samples (Supplementary Figure 4a,b). Notably, this representation failed to distinguish elongated and round negative-cell populations, which were resolved by CellMap. In contrast, CellMap embeddings yielded cell phenotypes that were consistently represented across all six samples (Supplementary Figure 4c), indicating greater robustness to sample-specific variation. We further evaluated the intensity-based representation in the 43-plex dataset (Supplementary Figure 4e). Because the baseline relies on marker intensity without explicitly encoding the spatial organization of the signal within the cell, it could classify cells as marker-positive when signal was detected outside the expected cellular compartment or originated from neighbouring cells. Representative examples include spurious or mislocalized marker signals that led to biologically implausible phenotype assignments (Supplementary Figure 4d).

#### Effect of clustering resolution

To assess the sensitivity of cell type discovery to the Leiden resolution parameter, we evaluated the 7-plex tonsil dataset across a range of clustering resolutions. As expected, increasing the resolution progressively increased the number of detected phenotypes, reflecting a finer subdivision of the cellular representation (Supplementary Figure 5A). To further assess whether this increased granularity altered the biological interpretation of the phenotypes, clusters obtained at each resolution were mapped to the seven major cellular classes identified at the reference resolution. The Sankey representation shows that increasing the resolution primarily subdivided existing biological populations into finer clusters, while preserving their correspondence with the original cellular classes (Supplementary Figure 5B). Consistently, the abundance of the major cellular classes at each resolution showed a strong correlation with their abundance at the reference resolution (0.3) across images (Supplementary Figure 5C), indicating that changes in clustering resolution predominantly affect phenotype granularity while preserving the underlying biological composition. We additionally evaluated the effect of the Leiden resolution parameter on neighborhood identification in the 43-plex tonsil dataset. Increasing the clustering resolution resulted in a progressively finer subdivision of tissue neighborhoods, while preserving the major spatial organization patterns identified at lower resolution (Supplementary Figure 6). These results indicate that, similarly to cellular phenotypes, the resolution parameter primarily controls the granularity of the inferred tissue organization without substantially altering its underlying spatial structure.

#### Quantitative evaluation of CellCut

CellCut was initially evaluated qualitatively by visual inspection of representative examples illustrating the correction of nucleus–cell pairing errors (Fig. 2). In addition, the nuclear segmentation component was independently evaluated against manual annotations, achieving an F1-score of 0.89. To quantitatively assess the contribution of the CellCut nucleus–cell pairing correction, we manually annotated 10 representative 100×100-pixel image crops from the 7-plex tonsil dataset. These crops were selected to cover a range of cell densities and spatial distributions. We evaluated whole-cell segmentation using the same DeepCell cell segmentation as the baseline, before and after application of the CellCut watershed-based nucleus–cell pairing correction. The baseline DeepCell segmentation yielded a mean F1-score of 0.68 ± 0.12, whereas the application of the CellCut correction improved performance to 0.75 ± 0.09.

## Discussion

The increasing availability of multiplex and hyperplex imaging technologies enables the study of tissue organization schemes at a single-cell resolution but also introduces substantial computational challenges for data analyses. In this work, we presented UMITIC, a fully unsupervised and modular framework that addresses the key limitations of the current multiplex imaging analysis techniques. Specifically, UMITIC combines accurate a paired nucleus–cytoplasm single-cell segmentation process through CellCut, a contrastive self-supervised representation learning scheme tailored to high-dimensional multiplex single-cell imaging through CellMap, cellular embeddings that integrate both phenotypic and morphologic information, and TissueNet, a lightweight graph neural network that was specifically developed to model spatial cell– cell interactions and infer tissue neighborhoods in a fully data-driven manner. Together, these components enable interpretable tissue characterization work to be performed across both cellular phenotypes and higher-order spatial organization schemes.

A central component of the proposed approach is its use of contrastive self-supervised learning to generate low-dimensional representations of multichannel single-cell images. This strategy enables the extraction of feature representations that capture biologically relevant features while achieving reduced dimensionality and mitigating the technical variability that arises from staining, imaging and intersample differences, thereby eliminating the need for prior preprocessing. By concatenating both marker expressions and morphological descriptors, the resulting embeddings capture complementary aspects of cellular identities that are often difficult to manually quantify. Importantly, cellular morphologies provide biologically relevant information that is associated with pathological and functional cellular states. For example, cell and nuclear morphology alterations have long been linked to cancer progression and tumor aggressiveness.

The inclusion of explicit cell segmentation through the CellCut module further enhances the biological interpretability of the framework. Beyond the qualitative improvements, the quantitative evaluation demonstrated that the CellCut nucleus–cell pairing correction improved whole-cell segmentation accuracy compared with DeepCell segmentation. Moreover, by ensuring accurate nucleus–cytoplasm pairing, CellCut enables the precise extraction of morphological features, which can be associated with functional or pathological states. This paired segmentation also readily enables per-marker nuclear-to-cytoplasmic intensity ratios, a straightforward extension that could capture translocation-dependent signaling activity (e.g., FoxP3, Ki67 and Pax5) at no additional segmentation cost. While this step increases the computational complexity level, it provides a more faithful representation of individual cells than patch-based approaches do, as these methods may mix signals derived from multiple cell types. The benchmark comparison between UMITIC (cell-based) and NaroNet (patch-based) conducted on our in-house MI dataset further supports this statement.

At the tissue level, the TissueNet module incorporates spatial context through a graph neural network, enabling the identification of cellular neighborhoods on the basis of learned cell–cell interactions. Unlike methods that rely on predefined spatial features or fixed adjacency assumptions, this approach enables the constructed model to learn the relative importance levels of spatial relationships directly from the input data. The combination of cell-level and neighborhood-level clustering processes provides a multiscale representation of the tissue organization scheme, yielding improved interpretability and facilitating comparisons across samples.

Across datasets with varying complexity levels, UMITIC consistently recovered known biological structures. In tonsil tissue, the framework identified canonical immune populations and reconstructed well-established anatomical regions, including germinal centers, mantle zones, and T-cell areas. Importantly, these structures were preserved across both the 7-plex and 43-plex datasets, indicating the robustness of UMITIC to increasing marker dimensionality. Interestingly, the higher-dimensional dataset enabled a finer cell population stratification procedure, illustrating how the clustering resolution can be adjusted to capture different levels of biological granularity. In contrast, spatial neighborhood identification remained stable across different datasets, suggesting that higher-order tissue organization schemes can be consistently recovered despite marker panel differences and independent of the number of markers included.

UMITIC showed improved performance compared with that of the existing state-of-the-art methods across both qualitative and quantitative assessments. Qualitatively, the comparison with NaroNet [20] in Fig. 4 underscores the superior ability of our method to reconstruct tissue architectures across multiple levels of tissue organization: the UMITIC single-cell type mosaics (Fig. 4d) showed identifiable, consistent and biologically relevant cell phenotypes in human tonsils, whereas the NaroNet patches (Fig. 4e), even when they were adjusted to single-cell size, showed less consistent cell types, often mixing various biologically relevant cell phenotypes in a single group. Moreover, while UMITIC was able to reconstruct the tonsil histology at the tissue neighborhood level with remarkable accuracy, recognizing and separating known tonsillar regions—B follicles, germinal centers, mantle, T-cell zones and the epithelium—in a completely unsupervised way and without the need for annotated labels, NaroNet yielded a more mixed picture when its patches were clustered at the cell neighborhood level (Fig. 4f). In addition, NaroNet relies on an MLP architecture that requires manually predefining a fixed number of tissue microenvironment elements, such as cell types or neighborhoods, before conducting training. This design constrains the flexibility of the model and may bias the learned representations toward those imposed categories. In contrast, UMITIC employs an unsupervised clustering strategy to identify both cellular phenotypes and spatial neighborhoods directly from the given data, allowing their number and composition to adaptively emerge. Quantitatively, we evaluated the performance of our TissueNet module on an external, annotated healthy lung IMC dataset, further demonstrating the adaptability of UMITIC across different tissue types and imaging modalities. We compared it with UTAG [19] and CytoCommunity [10], state-of-the-art reference methods for the unsupervised characterization of tissues from MI via the definition of spatial neighborhoods. Notably, this evaluation conducted using an expert-annotated lung dataset confirmed that UMITIC accurately delineated the principal anatomical compartments of the lung, significantly outperforming both UTAG and CytoCommunity in terms of homogeneity and Rand index scores (p < 0.001, paired Wilcoxon signed-rank test). A visual assessment of the tissue samples (Fig. 7a) revealed the remarkable accuracy achieved by our method in comparison with that of UTAG with respect to the reconstruction of distinct lung microanatomical compartments.

The generalizability of our method was further supported by an experiment conducted on an external 58-plex colorectal cancer cohort [23], where UMITIC recovered previously reported immune composition and spatial organization differences between two patient groups involving different prognoses. Remarkably, all the group comparison results, both for the cell types and tissue neighborhoods of interest, found by UMITIC were consistent with those reported by the authors of the original paper [37]. The minor discrepancies observed in neighborhood abundances may be explained by differences in spatial resolution, as UMITIC performs the analysis at single-cell resolution, whereas the original study used a sliding-window approach that aggregates information from multiple cells. Consequently, our approach can capture fine-grained biological heterogeneity and subtle local structures that may be smoothed by window-based spatial aggregation. Conversely, this higher spatial resolution also means that segmentation errors during the CellCut step may have a greater impact on downstream neighborhood estimates.

Despite these strengths, the limitations of our approach should also be considered. First, the accuracy of downstream analyses depends on the quality of the initial cell segmentation results, and errors at this stage may propagate through the pipeline. However, CellCut is based on a dual-segmentation strategy that takes advantage of two state-of-the-art cell segmentation reference methods, postprocessing their segmentations to provide a final, accurate, paired nucleus cytoplasm single-cell delineation procedure. Moreover, we fine-tuned and tested the nuclear segmentation approach on an in-house database, achieving high segmentation accuracy with an F1 score of 0.89, and quantitatively demonstrated that our pairing correction improves overall whole-cell segmentation accuracy. Second, the identification of cell types and spatial neighborhoods relies on clustering, which introduces sensitivity to the Leiden clustering resolution parameter. While this provides flexibility, the resolution must be selected according to the desired level of biological detail, as higher values produce finer-grained structures and lower values yield broader groupings, thereby directly affecting the granularity of the obtained results. Third, the low availability of publicly accessible hyperplex datasets limits the quantitative evaluation of the pipeline, particularly in hyperplex datasets with more than 20 biomarkers per sample.

Taken together, our results support the hypothesis that unbiased, unsupervised characterizations of individual cells and their spatial organization schemes, particularly within tumor microenvironments, may yield novel and clinically relevant insights. By profiling cell types and their spatial distributions without imposing prior assumptions regarding their numbers or identities, our approach is designed to generalize effectively across diverse settings and to uncover latent patterns that might otherwise remain undetected. Future work will be focused on integrating these spatial and phenotypic features with patient-level clinical data, including treatment responses, disease progression trends, and tumor stages, through weakly supervised learning frameworks. This integration strategy could further advance precision oncology by enabling the identification of robust predictive biomarkers and supporting the development of personalized therapeutic strategies. This is particularly relevant in the context of immune checkpoint inhibitors, which currently benefit only a limited subset of patients, underscoring the urgent need for more reliable predictive markers [40]. Further extensions of this work may include the incorporation of additional data modalities, such as spatial transcriptomics, as well as model interpretability enhancements for better elucidating the contributions of individual cellular markers and morphological features. The modular architecture of UMITIC facilitates these extensions and ensures its adaptability to a wide range of experimental and clinical settings.

In summary, UMITIC provides a scalable and highly interpretable framework for the unsupervised analysis of multiplex and hyperplex imaging data, enabling the joint characterization of cellular phenotypes and their spatial organization schemes in tissue. By integrating molecular, morphological and spatial information, the proposed approach contributes to the development of reproducible and data-driven tools for studying tissue architectures in complex biological systems.

### Computational environment and reproducibility

All analyses were performed in Python 3.8.20 using TensorFlow 2.8.4, PyTorch 1.12.1, Keras 2.8.0, Scanpy 1.9.8, StarDist 0.9.1, DeepCell 0.12.10, PyTorch Geometric 2.1.0, and Leiden clustering (leidenalg 0.10.2). Deep learning models were trained and evaluated using an NVIDIA RTX A6000 GPU with CUDA 11.2 and cuDNN 8.1. The complete source code and configuration files required to reproduce the analyses are publicly available at https://github.com/mariasanguesa/UMITIC.

## Supporting information captions

**Supplementary Figure 1. UMITIC results derived from six human tonsil samples analyzed with a 7-plex multiplex panel**. **a.** Tissue reconstructions obtained based on cell type assignments. Top: original multiplex image crops; bottom: segmented cells colored according to the assigned cell type (T1–T8). **b.** Tissue reconstructions obtained by neighborhood (NB1–NB5) across five tonsil samples. **c.** Bar plot of the cell type composition per neighborhood

**Supplementary Figure 2. NaroNet results obtained for six human tonsil samples analyzed with a 7-plex multiplex panel**. a. Heatmap of the average marker expression per identified phenotype (Ph1–Ph10), normalized by marker. The cell counts are shown in green. **b.** Heatmap of the average marker expression per neighborhood (Nb1-Nb7), normalized by marker. The cell counts are shown in green.

**Supplementary Figure 3**. **Heatmap showing the mean per phenotype expressions of 58 protein markers**. The values are min–max normalized per marker (0–100), and phenotypes T01–T53 are named according to their marker profiles. The data are derived from an external 58-plex colorectal cancer cohort.

**Supplementary Figure 4. Comparison of CNN-based CellMap embeddings and per-cell marker-intensity representations for cell type identification. a.** Heatmap of the mean marker-expression profiles of cell types identified in the 7-plex tonsil dataset using per-cell marker-intensity vectors. **b**. Relative abundance of the phenotypes identified from marker-intensity vectors across the six tonsil samples. **c.** Relative abundance of the cell types identified using CellMap embeddings across the six samples. **d**. Representative examples illustrating phenotype assignments obtained from marker-intensity vectors and CellMap embeddings in the 43-plex tonsil dataset. Mislocalized or spatially ambiguous marker signals can result in positive assignments when only marker intensity is considered. **e.** Heatmap of the mean marker-expression profiles of cell types identified in the 43-plex dataset using per-cell marker-intensity vectors.

**Supplementary Figure 5. Effect of the Leiden clustering resolution on phenotype granularity and biological composition in the 7-plex tonsil dataset. a.** Number of cell types identified across different Leiden resolution values. **b.** Sankey diagram showing the correspondence between the seven major biological cell classes identified at the reference resolution (0.3) and the finer phenotypic clusters obtained at resolution 0.7. **c.** Correlation between the abundance of the major cell classes at the reference resolution (0.3) and at increasing clustering resolutions across the six tonsil samples. Each point represents one image; Pearson correlation coefficients are indicated in each panel.

**Supplementary Figure 6. Effect of Leiden clustering resolution on tissue neighborhood granularity in a 43-plex tonsil image.** Tissue neighborhood assignments obtained from the same 43-plex tonsil image using increasing Leiden resolution values (0.3, 0.5, 0.7, 0.9 and 1.2). Higher resolutions progressively subdivide the inferred neighborhoods while preserving the major spatial organization of the tissue.

