## Supplementary figures and images for "UMITIC: An unsupervised framework for the joint characterization of cellular phenotypes and spatial neighborhoods in multiplex and hyperplex immunofluorescence imaging data"

### Supplementary Figure 1

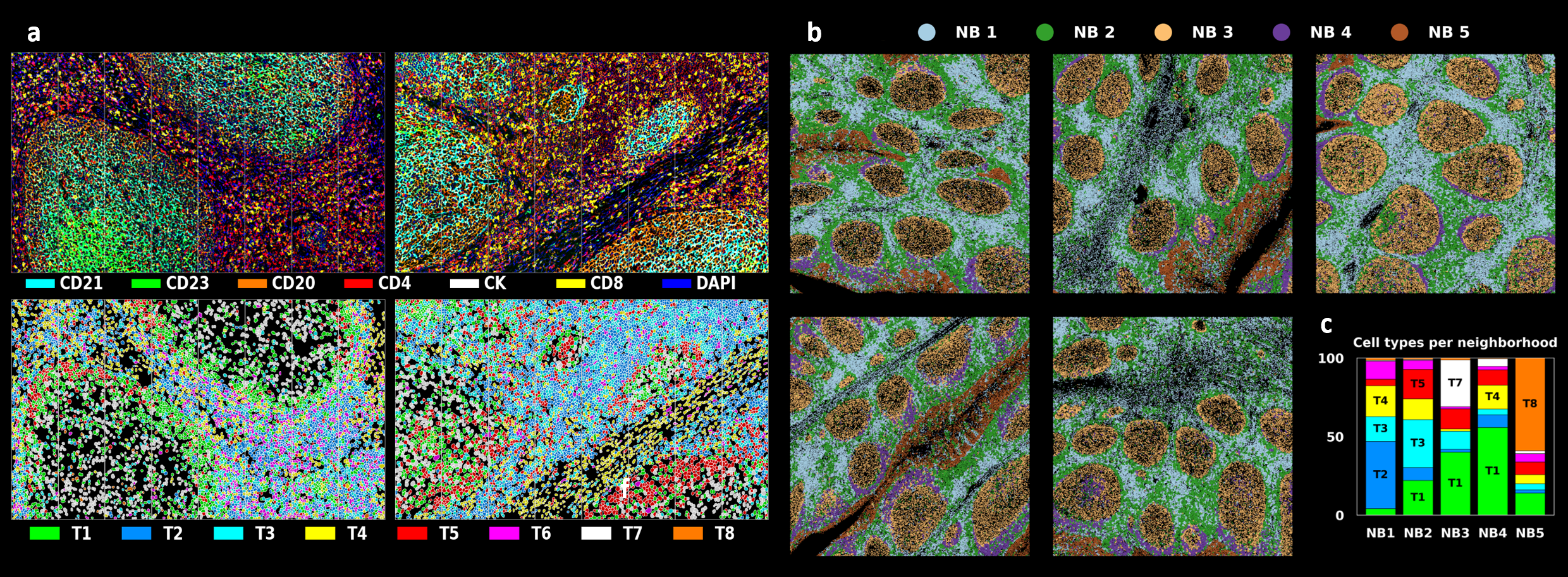

### Supplementary Figure 2

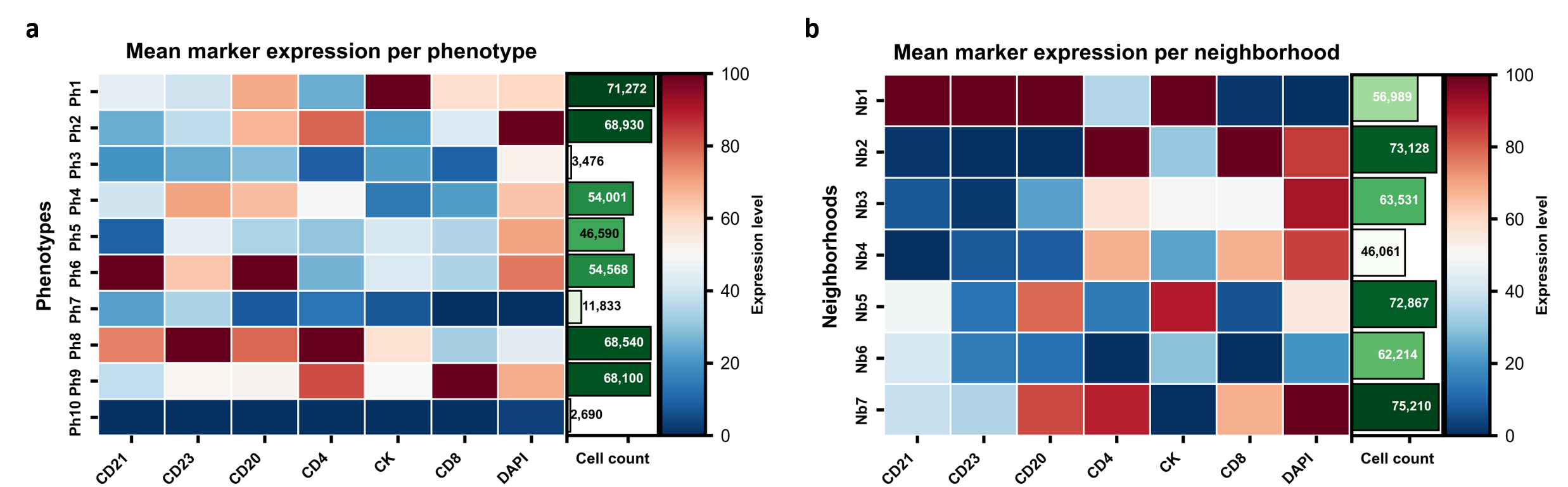

### Supplementary Figure 3

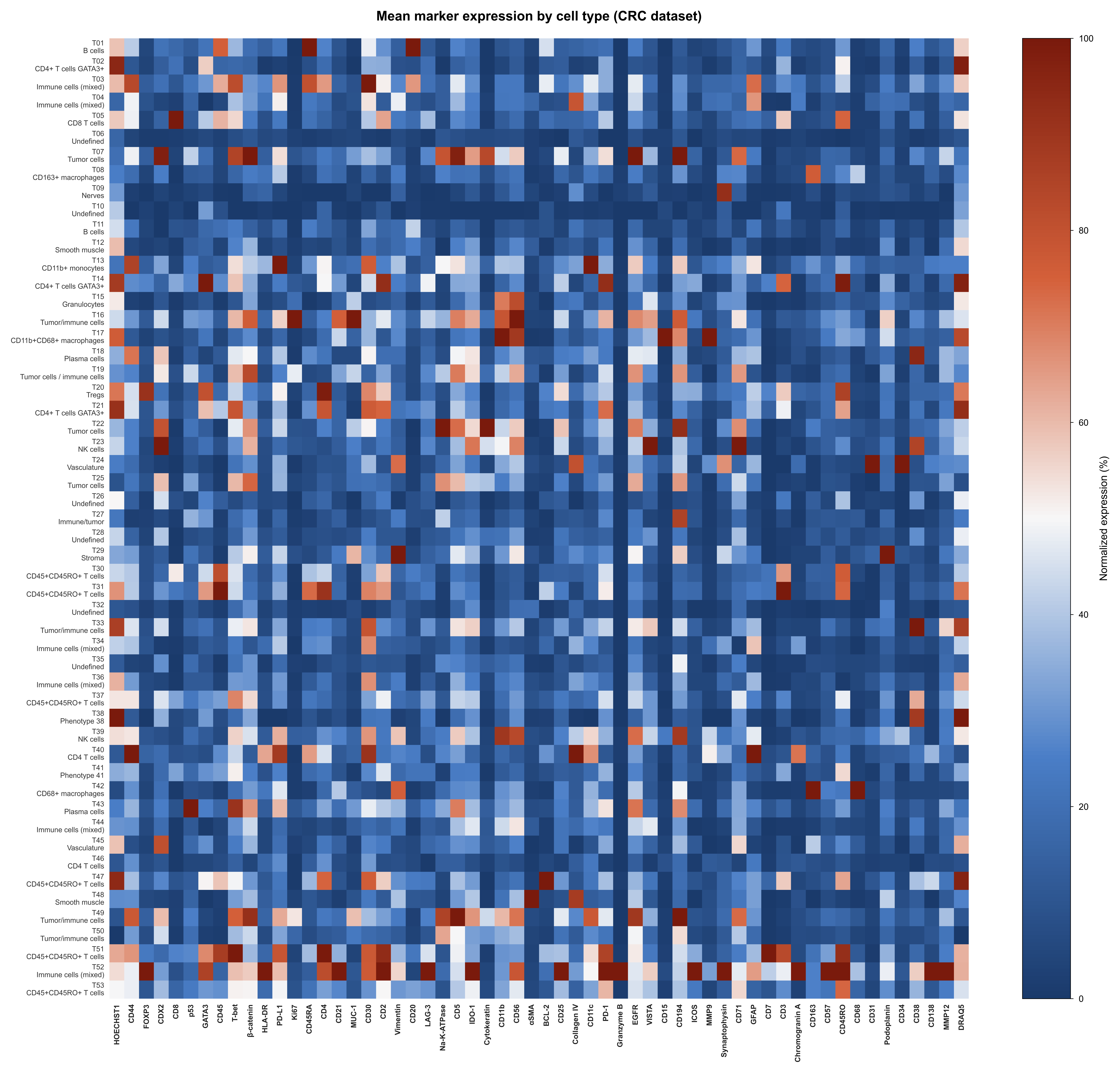

### Supplementary Figure 4

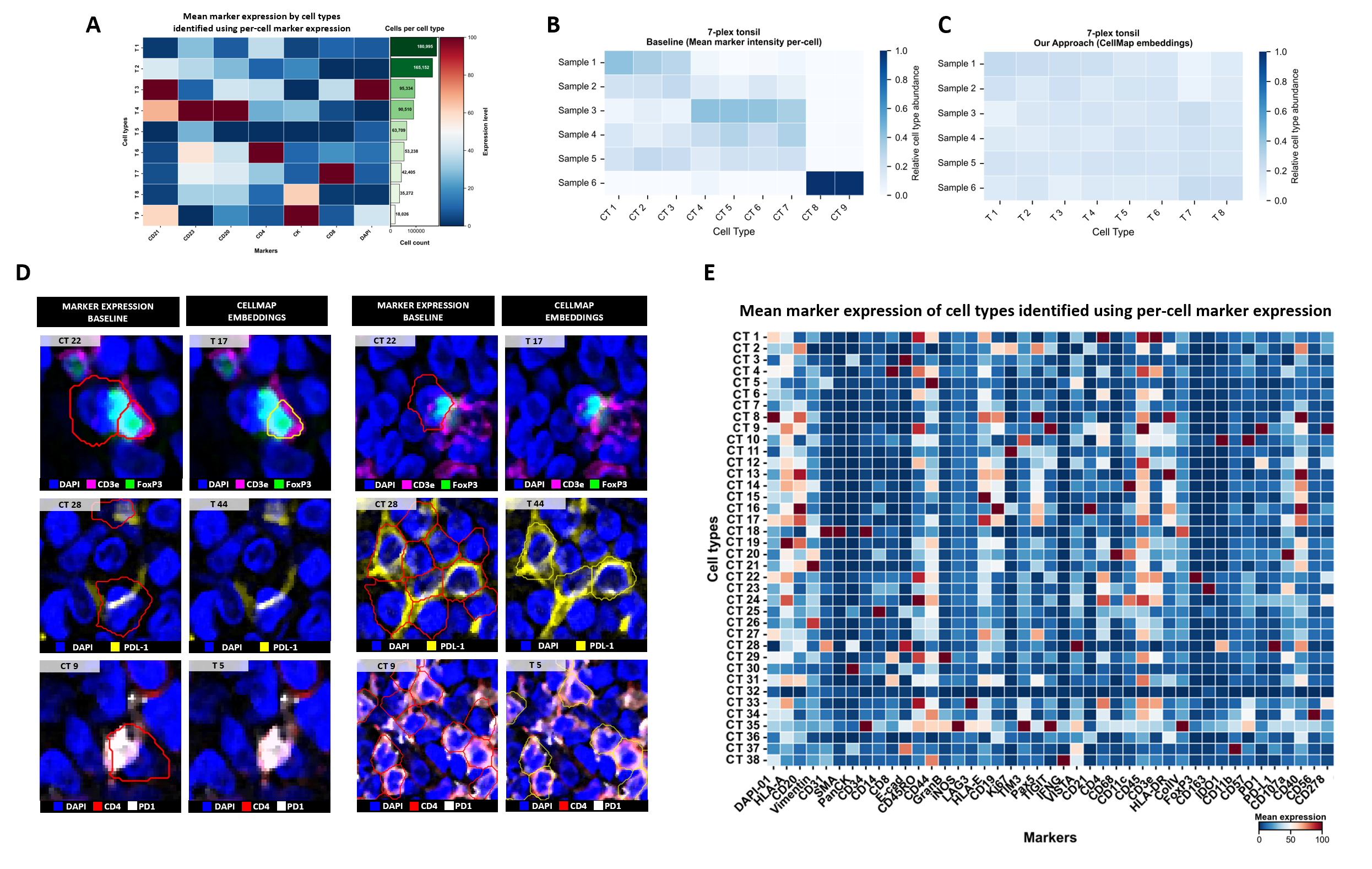

### Supplementary Figure 5

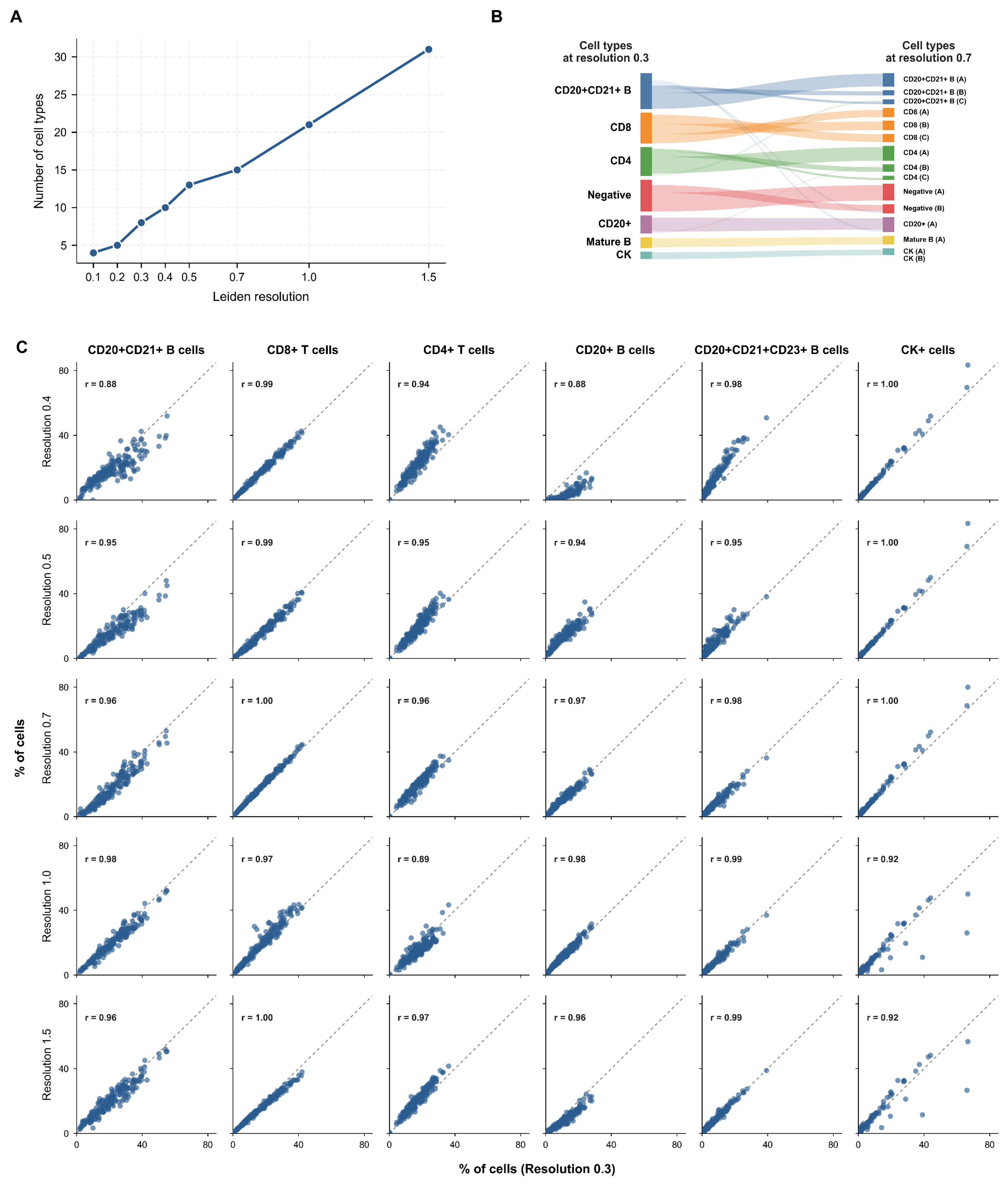

### Supplementary Figure 6

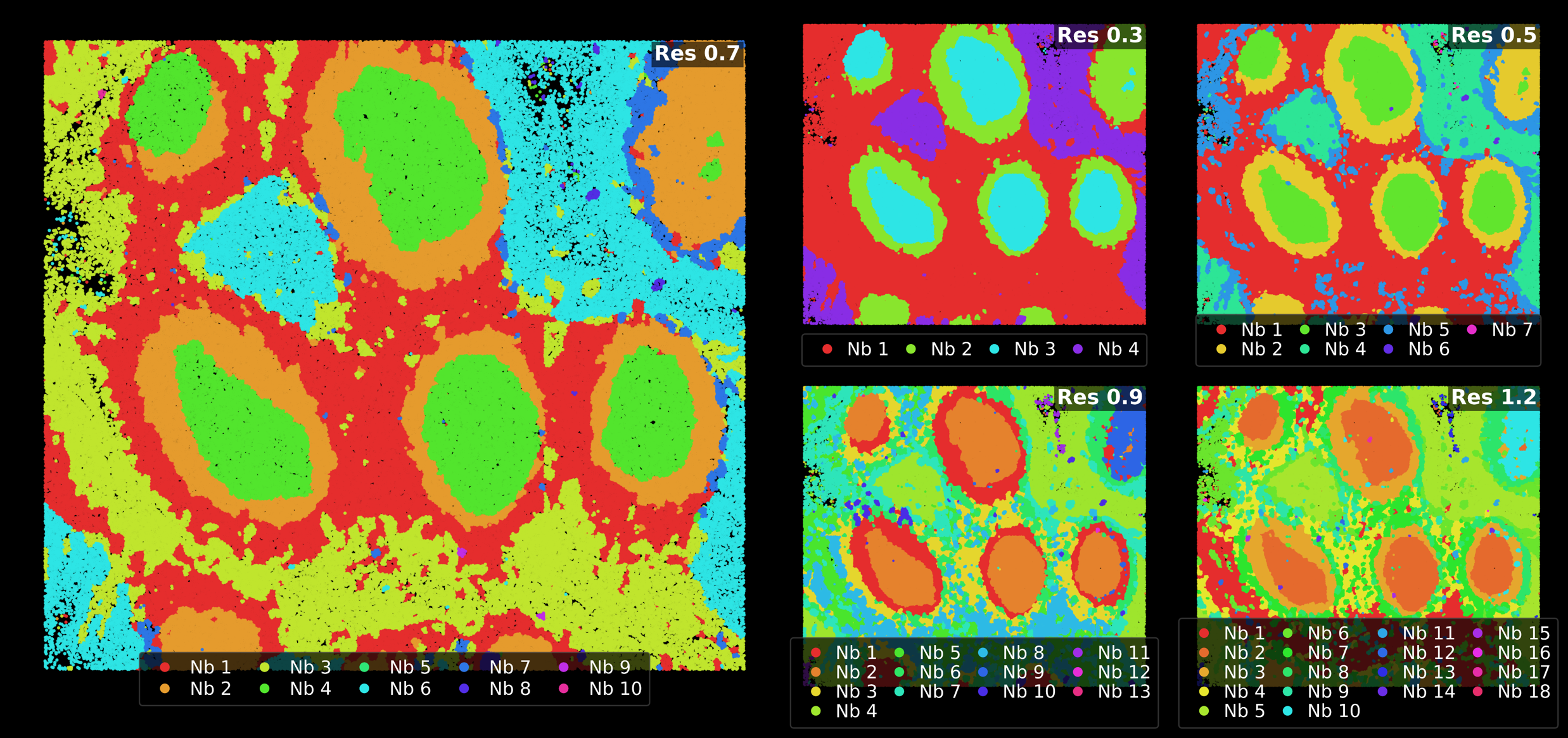
